# Coaxially Electrospun Myocardial dECM- based Nanofibrous Scaffolds Demonstrate Enhanced Cardiomyocyte Adhesion and Function

**DOI:** 10.1101/2025.09.02.673432

**Authors:** Dhanusha N. Rajapakse, Kiran M. Ali, Mahtab Khodadadi, Bryson T. Proctor, Tanya Upadhyay, Daxian Zha, Taylor C. Suh, Jessica M. Gluck

## Abstract

Limited regenerative capability of mature cardiomyocytes (CMs) makes myocardial repair more challenging, requiring effective and viable alternatives to conventional heart transplants. Cardiac tissue engineering is a substitutionary approach combining cells, scaffolds, and growth factors to develop functional heart tissues in vitro. Induced pluripotent stem cells (iPSCs) represent a significant advancement in cardiac regenerative medicine, offering a continuous supply of CMs, however, the limited understanding of their microenvironment hinders translational research. Decellularized extracellular matrix (dECM) derived from myocardium is a highly promising natural scaffold for CTE, given its tissue-specific composition, mechanical properties, and biochemical cues that promote cellular regeneration. This study investigates myocardial dECM-based fibrous scaffolds for iPSC-derived CM use. Coaxially electrospun nanofibers comprising a polyurethane core and a blend of polycaprolactone and myocardial dECM as the sheath were optimized. Morphological analysis confirms the resemblance of the nanofibers to fibrillar collagen in the native dECM. ATR-FTIR and immunostaining results confirm the presence of dECM which enhanced their hydrophilicity and enzymatic degradation. Biocompatibility results show higher phenotypic retention of iPSC-CMs due to microenvironments enriched with native proteins. On the contrary, the scaffolds without myocardial proteins exhibit higher dedifferentiation of iPSC-CMs, proving that ECM proteins provide a suitable microenvironment for iPSC-CMs.

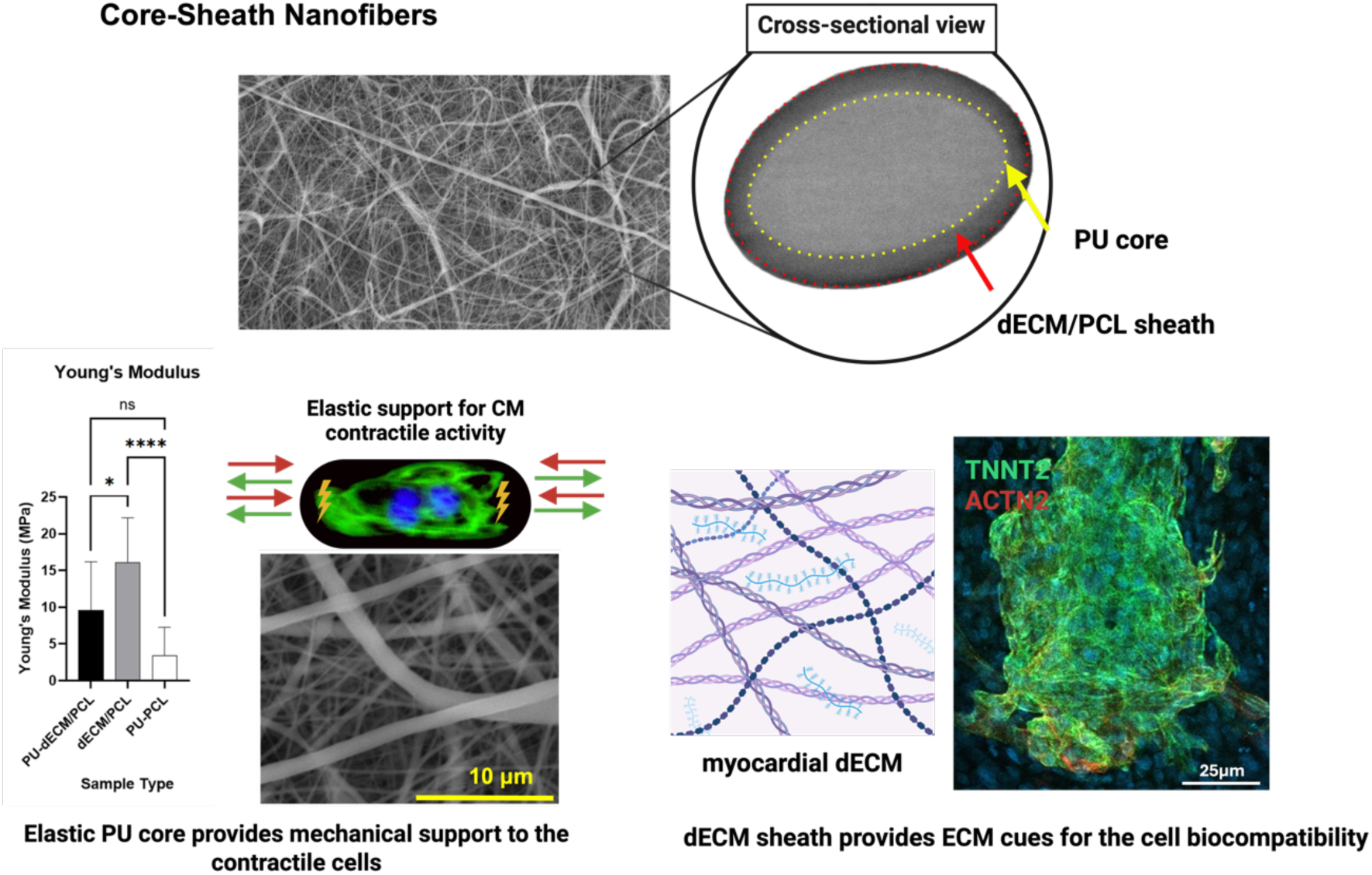

## 1. Introduction

Cardiac tissue engineering (CTE) is dedicated to using tissue engineering principles to develop contractile and functional cardiac tissues in vitro for in vivo regenerative purposes.^[1]^ The discovery of induced pluripotent stem cells (iPSCs) has been pivotal, enabling the creation of customized human tissues from somatic cells.^[2]^ Cardiomyocytes (CM) are the contractile cells of the heart that are prominently present in the heart walls, crucial for the beating function of the heart. CMs exhibit a limited regenerative capacity, often being replaced by stiff scar tissue that impairs their function. Beyond tissue regeneration, CTE also supports heart disease modeling, facilitating the study of pathophysiology, and in vitro testing for drug safety and efficacy^[3–5]^.

Traditional CTE approaches use biocompatible scaffolds for delivering cells and growth factors to support subsequent tissue repair and functional restoration^[1–3,6,7]^. Ideally, a scaffold for CTE should match the structural, compositional, and functional properties of the native cardiac ECM. In addition to that, the scaffold should possess adequate strength and elasticity to support and withstand the systolic and diastolic functions of the heart^[7,8]^.

Electrospinning is a technique that produces micro or nanoscale fibers from a polymeric melt or solution, under a high electric field^[7,9,10]^. It is the most preferred method to make scaffolds for CTE due to its simplicity, versatility, comparatively less material restrictions, cost-effectiveness, and better control over the size and morphology of the fibers and porosity of the scaffold. It has established its success in producing biocompatible nanofibrous scaffolds that can closely mimic the ECM owing to its ability to generate fibers with sizes comparable to those of the fibrous ECM proteins such as collagen, elastin, and fibronectin^[10–12]^.

At present, a number of derivatives or modified versions of the traditional electrospinning process have been investigated for fabricating nanofibers. Coaxial electrospinning is one such novel electrospinning type which enables the production of fibers with a core-sheath structure in a single step^[13–15]^. This technique utilizes a concentric set of capillaries that supports ejecting two or more solutions simultaneously, where the smaller needle feeds the core solution and the larger needle supplies the shell solution^[9,14]^. These co-flowing solutions form a compound jet at the capillary tip, resulting in composite fibers^[16]^. The biggest advantage of this technique is that it allows the fabrication of complex multi-constituent fibers and thereby achieves a synergy between different properties unattainable with a single material^[15]^. It addresses challenges like miscibility and cross-reactivity when blending different materials and allows for the incorporation of typically insoluble substances within the core. The structural arrangements from this technique are not limited to single core-sheath structures and include hollow fibers, triple/quadruple layered fibers, and multi-channeled fibers^[15]^. In addition to that, coaxial electrospinning also favors fiber-based drug carriers, as it enables the controlled release of drugs and growth factors ^[9,16]^.

Both natural and synthetic materials can be used for the fabrication of CTE scaffolds.^[17,18]^ Studies have shown that the use of composite scaffolds in TE has yielded better results compared to scaffolds made of individual counterparts^[5]^. Polycaprolactone (PCL) and polyurethane (PU) are two synthetic biopolymers that are being used in already commercialized FDA approved biomedical products^[3]^. Owing to their excellent biocompatibility and mechanical properties, they are among the most widely used synthetic biopolymers in biomedical applications including TE^[19,20]^. Both these synthetic biopolymers have been used in many tissue engineering applications that targeted different tissues and organs such as the heart, skin, cornea, liver, nerves, blood vessels, and bones^[21]^.

dECM from myocardium is extensively studied for its potential as a natural biological scaffold in cardiac tissue engineering. Its organ-specific ECM microstructure and composition, tissue-mimetic mechanical properties, and site-specific biochemical cues promote cardiac tissue regeneration^[8,22]^. As a tissue-derived dECM type, it retains tissue-specific memory, aiding the tissue-specific differentiation^[8]^. Proteomic studies reveal that decellularized human and porcine myocardium tissues consist of a complex mixture of around 200 cardiac ECM proteins such as collagen, elastin, laminin, and glycosaminoglycans that are vital for cell attachment, proliferation, and differentiation^[22]^. In addition to that, the presence of soluble matrix-bound growth factors and cytokines has also been detected in those decellularized tissues^[22]^.

Deng et al. reported on coaxially electrospun aligned nanofibers for nerve tissue regeneration, featuring a PCL core and dECM from peripheral nerves in the sheath^[23]^. While coaxial electrospinning shows promise for dECM-based scaffolds in tissue engineering, its application for cardiac-derived scaffolds remain underexplored. This study addresses that gap by creating dECM-based core-sheath, characterizing their properties and evaluating biocompatibility to support the differentiation of iPSC-CM in vitro. We developed a bioactive, mechanically robust scaffold made of core-sheath bicomponent fibers using coaxial electrospinning where polyurethane serves as the elastic core and a blend of PCL with porcine myocardial dECM forms the sheath. Incorporating dECM into the sheath aims to enhance biocompatibility, while dual synthetic polymers improve the mechanical properties.

## 2. Results

### 2.1. Coaxial electrospinning of the three electrospun nanofiber types and their structural and morphological analyses

Bicomponent nanofibers with core-sheath configuration were fabricated using coaxial electrospinning where a solution containing a mixture of PCL and porcine myocardial dECM formed the sheath and a solution of PU formed the core. To compare their physicochemical, and functional properties, two control sample types were determined - (a) monoaxially electrospun monolithic fibers containing a blend of PCL and dECM (dECM/PCL) and (b) coaxially electrospun bicomponent fibers with a PU core and PCL sheath (PU-PCL). The purpose of having dECM/PCL fibers as a control was to study the changes in properties and performance brought about by changing the fiber configuration from monolithic to bicomponent. Meanwhile, the PU-PCL control was included to investigate the variations in properties and performance resulting from the presence and absence of dECM, despite having the same fiber architecture. **Table 1** provides a summary of the three different fiber types, their method of fabrication and composition.

**Table 1.**
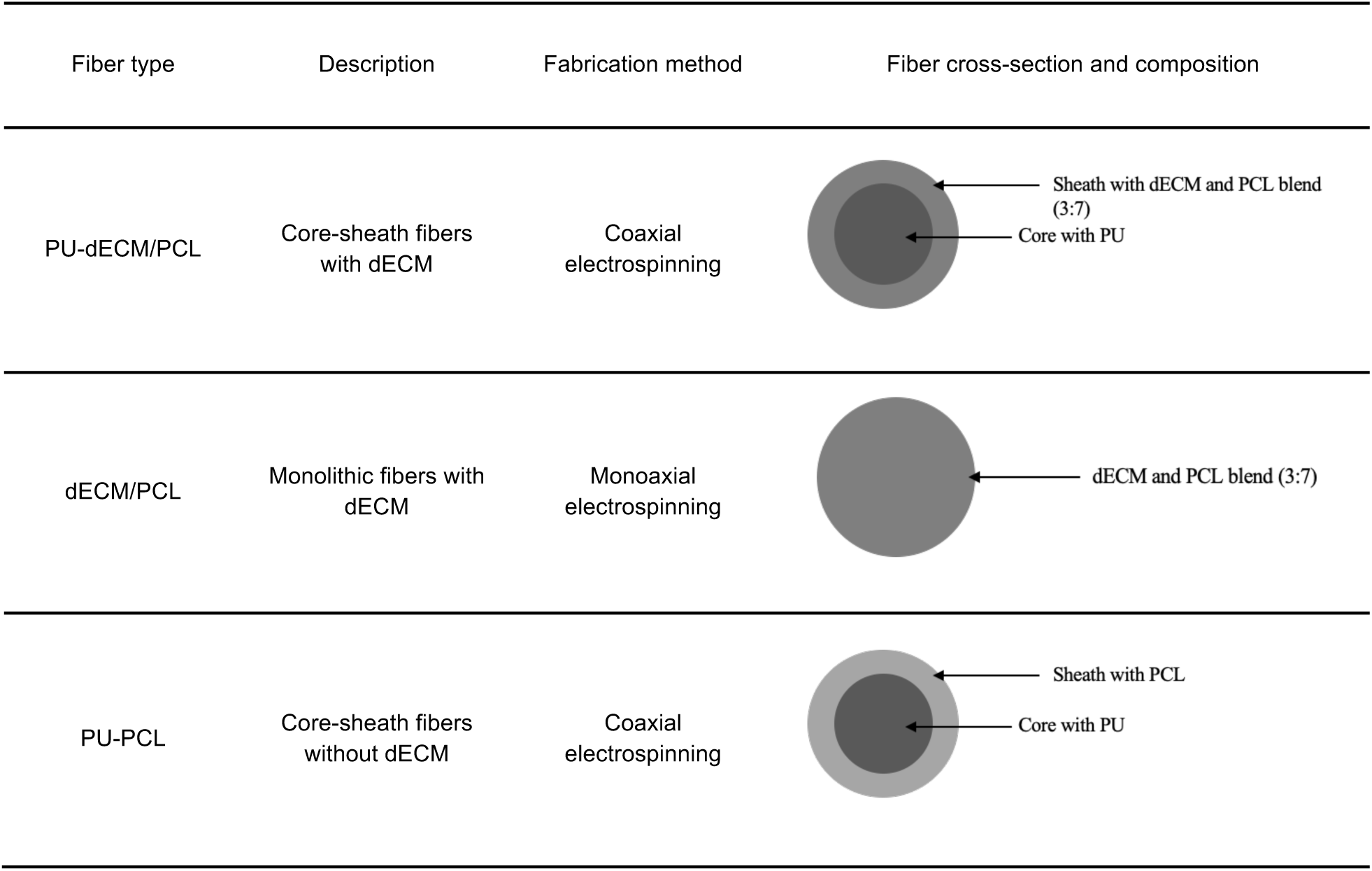
Summary of the fabricated fiber types.

| Fiber type | Description | Fabrication method | Fiber cross-section and composition |
| --- | --- | --- | --- |
| PU-dECM/PCL | Core-sheath fibers with dECM | Coaxial electrospinning | Sheath with dECM and PCL blend (3:7)<br>Core with PU |
| dECM/PCL | Monolithic fibers with dECM | Monoaxial electrospinning | dECM and PCL blend (3:7) |
| PU-PCL | Core-sheath fibers without dECM | Coaxial electrospinning | Sheath with PCL<br>Core with PU |

TEM images of the PU-dECM/PCL fiber cross-sections (**Figure 1a**) show distinctive inner and outer regions with a clear boundary in between as opposed to those of dECM/PCL fibers (**Figure 1b**) that showed an even shading throughout the cross-section. These results confirmed the successful formation of the core-sheath fiber architecture via coaxial electrospinning. Such distinct core-sheath regions were observed in the majority of the fibers in the evaluated PU-dECM/PCL samples (**Figure S1**).

**Figure 1.**
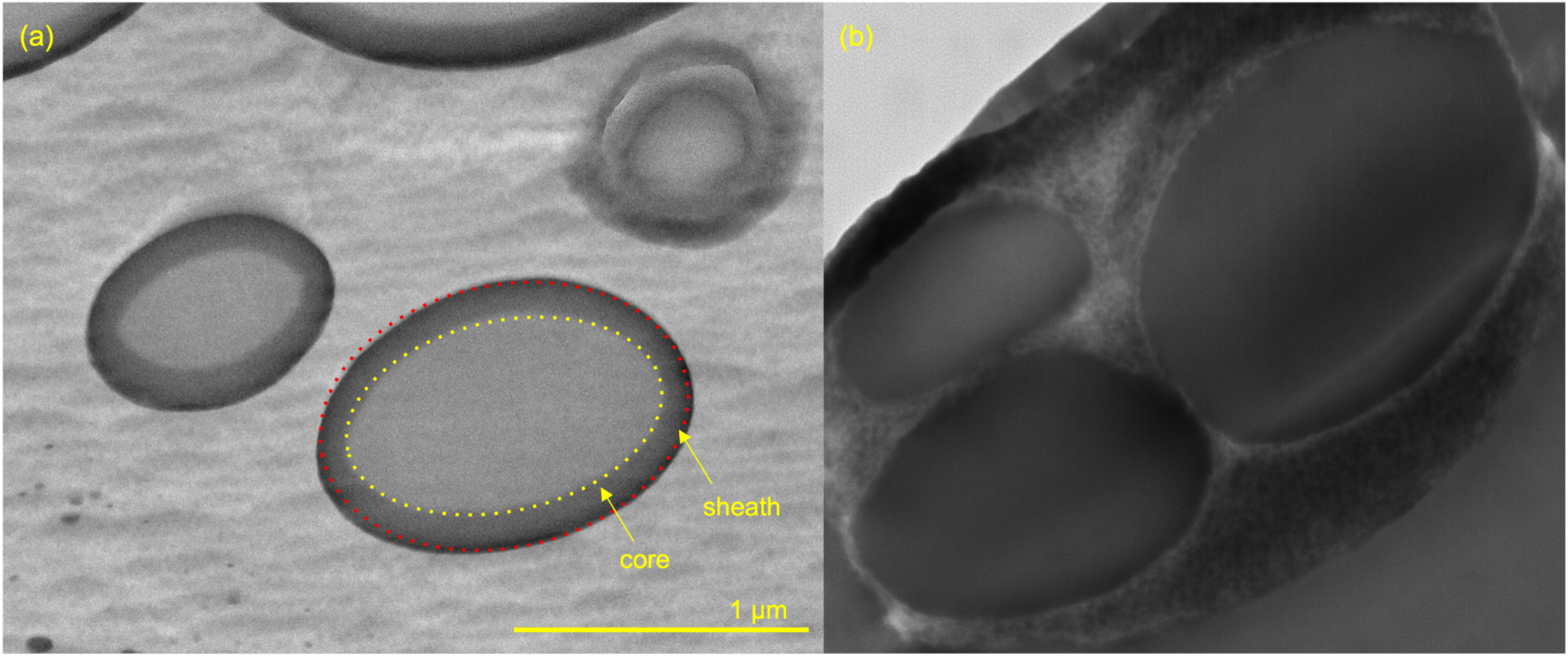
Confirmation of the presence of core-sheath structure in PU-dECM/PCL fibers. Cross-sectional TEM Images of (A) coaxially electrospun PU-dECM/PCL fibers, (B) uniaxially electrospun dECM/PCL fibers at 20,000x magnification. The red dotted lines indicate the outer circumference (sheath) of the fiber, while the yellow dotted line indicates the inner circumference (core). Scale bar = 1μm

SEM micrographs of the three fiber sample types (**Figure 2**) confirmed their uniform shape and morphology and the random arrangement without any identifiable orientation. The calculated values of average fiber diameter and pore size values for each sample type are given in **Table 2** and the respective distributions are given in **Figure 2**. The formation of continuous fibers without any fragments confirmed the use of optimum electrospinning parameters during the fabrication process. Continuous fibers with an even morphology provide a consistent substrate for the cells to attach and grow. Moreover, they are crucial for maintaining the mechanical integrity of the substrate.

**Figure 2.**
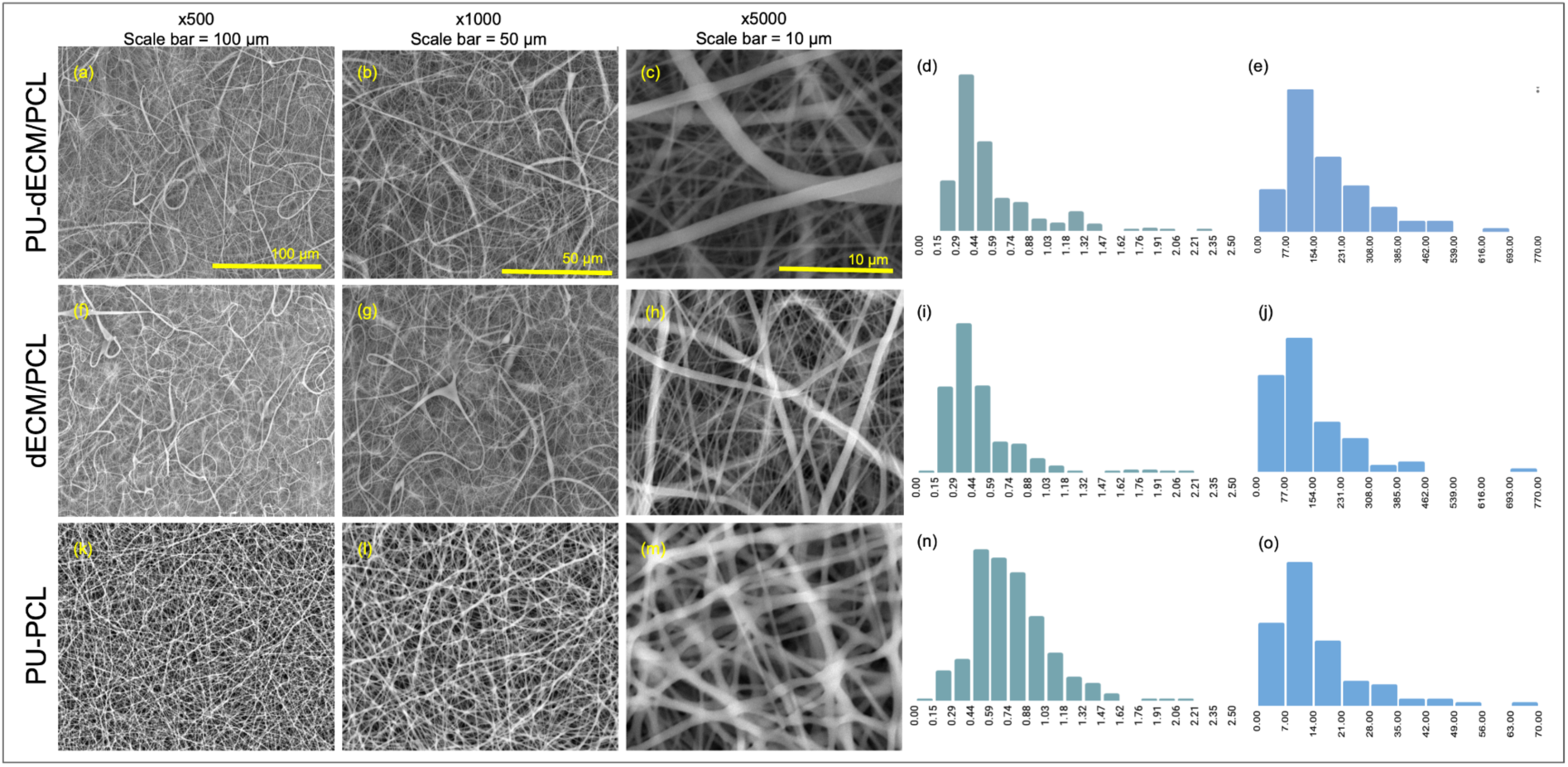
Size and morphological analysis of the three electrospun fiber types. SEM images PU-dECM/PCL fibers at (a) x500 (b) x1000 and (c) x5000 magnifications, dECM/PCL fibers at (f) x500 (g) x1000 and (h) x5000 magnifications, PU-PCL fibers at (k) x500 (l) x1000 and (m) x5000 magnifications, fiber diameter distribution of (d) PU-dECM/PCL fibers, (i) dECM/PCL fibers, and (n) PU-PCL fibers, pore size distribution of (e) PU-dECM/PCL fibers, (j) dECM/PCL fibers, and (0) PU-PCL fibers

**Table 2.**
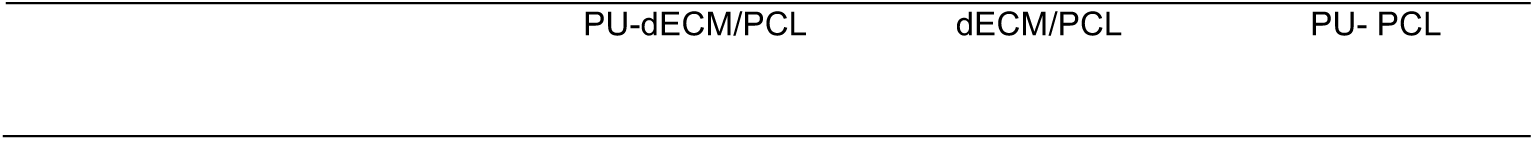
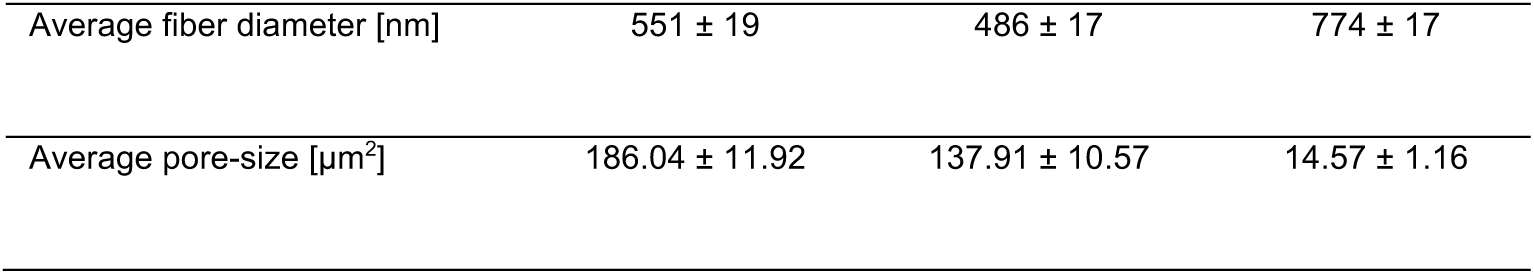
Calculated average fiber diameter and pore size of PU-dECM/PCL, dECM/PCL, and PU-PCL nanofibers (mean ± standard error of the mean).

| PU-dECM/PCL | dECM/PCL | PU- PCL |
| --- | --- | --- |

|  |  |  |  |
| --- | --- | --- | --- |
| Average fiber diameter [nm] | 551 ± 19 | 486 ± 17 | 774 ± 17 |
| Average pore-size [$\mu\text{m}^2$] | 186.04 ± 11.92 | 137.91 ± 10.57 | 14.57 ± 1.16 |

Morphological variations (such as beads) were visible in the SEM images of both PU-dECM/PCL (**Figure 2a**) and dECM/PCL (**Figure 2f**) fibers, as opposed to those of the PU-PCL (**Figure 2k**) fibers which had a clean appearance free of any beads or distortions. PU-PCL recorded the highest average fiber diameter followed by PU-dECM/PCL and dECM/PCL. Both core-sheath and monolith fibers containing dECM had similar fiber diameter distributions with most fibers belonging to the 290 – 440 nm range. The average fiber diameter of the core-sheath fibers with dECM was slightly higher and statistically significant than that of the monolith fibers (*p = 0.012). However, on average, core-sheath fibers with dECM were significantly smaller (***p < 0.001) as opposed to those without.

PU-dECM/PCL and dECM/PCL fibers had comparable average pore sizes (186.04 ± 11.92 μm² and 137.91 ± 10.57 μm², respectively, **Table 2**), though significantly different (**p = 0.003). Their pore-size distributions were also comparable with most pores in the 30–125 μm² range. In contrast, PU-PCL fibers had significantly smaller pores (14.57 ± 1.16 μm², ***p < 0.001). Accordingly, core-sheath fibers with dECM created the most porous substrate followed by monolithic fibers and core-sheath fibers without dECM.

### 2.2. Compositional analysis of the three electrospun nanofiber types

#### 2.2.1. ATR-FTIR analysis

ATR FT-IR analysis was carried out to verify the composition of the sheath of PU-dECM/PCL fibers, specifically to confirm the presence of dECM and PCL blend and the absence of PU on the sheath. ATR FT-IR spectra of the individual constituents and the PU-dECM/PCL fibers obtained at the wavelength range 4000-600 cm^−1^ (**Figure 3**).

**Figure 3.**
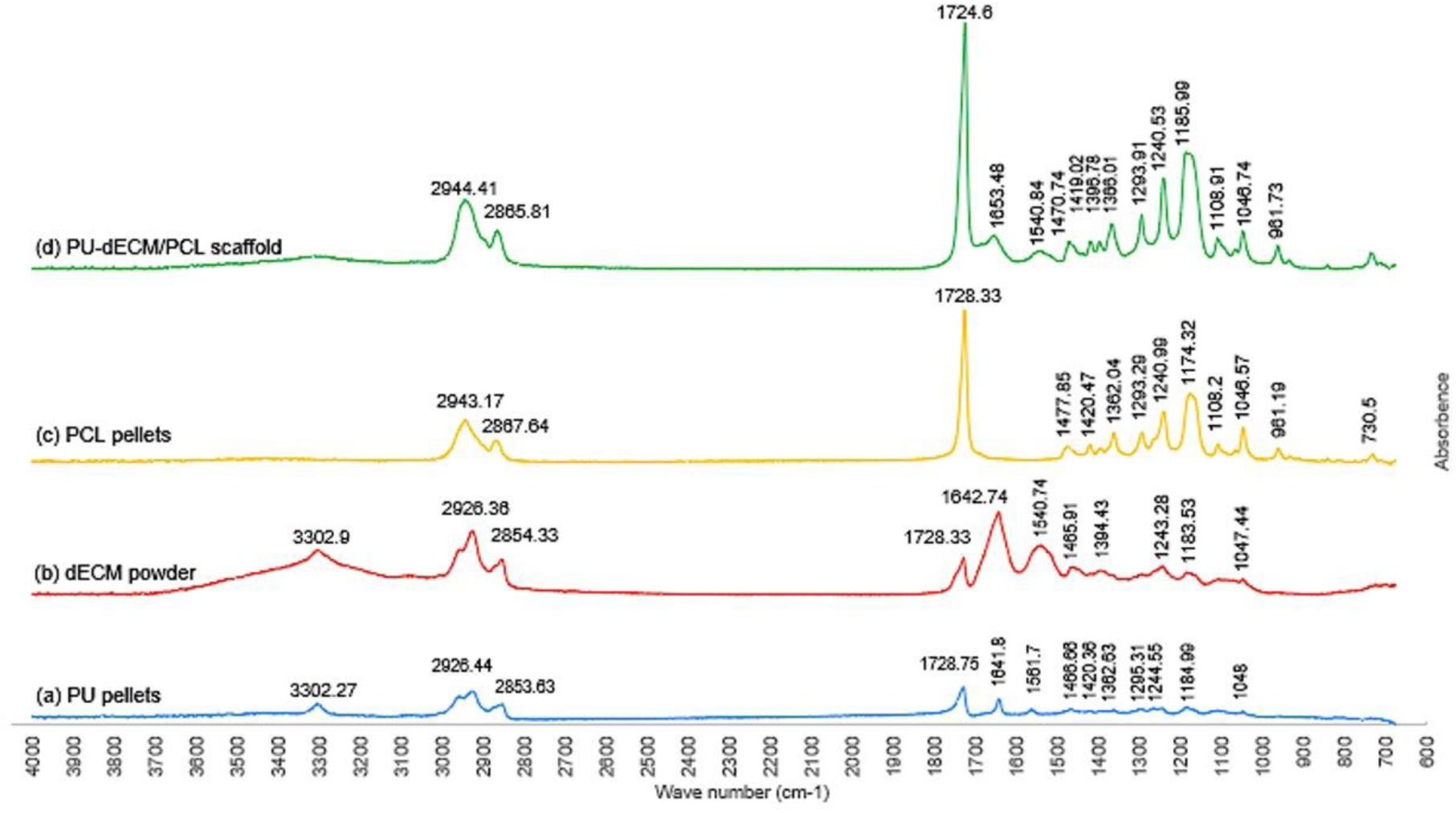
Surface chemical analysis of PU-dECM/PCL fibers. ATR FT-IR spectra of the (a) PU-dECM/PCL nanofibers - green (b) PCL pellets - yellow (c) dECM powder - red and (d) PU pellets - blue obtained at 4000 - 600 cm^−1^

In the PCL spectra (**Figure 3c**), the below characteristics were observed and were identified. The peaks seen at 2943 cm^−1^ and 2867 cm^−1^ corresponded to the asymmetric and symmetric stretching of C–H hydroxyl groups respectively and the peak at 1725 cm^−1^ represented the C– O stretching vibrations of the ester carbonyl group.^[24–26]^ The stretching and bending vibrations of –CH_2_ were represented by the peaks at 1477 cm^−1^ and 1420 cm^−1^ respectively.^[25–27]^ The peak at 1362 cm^−1^ corresponded to the stretching of the –OH group and the peak at 1293 cm^−1^ represented the stretching of the C–O bond.^[25–28]^ The asymmetric and symmetric stretching of the C–O–C bonds were shown by the peaks at 1240 and 1174 cm^−1^ respectively and the stretching of the oxime bonds was represented by the peak at 961 cm^−1^.^[24–26,28]^ All these peaks were observed in the FT-IR spectrum of the PU-dECM/PCL (**Figure 3d**) fibers at approximately equal wavenumbers. This confirmed the presence of PCL on the fiber surface. The two additional peaks observed at 1653 and 1540 cm^−1^ in the spectrum of the PU-dECM/PCL fibers corresponded to the stretching of C=O in amide groups and N–H bending vibrations respectively.^[26,29]^ These peaks were present on the spectra of both dECM powder (**Figure 3b**) and PU pellets (**Figure 3a**). Interestingly, the peak at 3302 cm^−1^ representing the stretching vibrations of N–H groups^[26,28,29]^ recorded in the individual spectra of both dECM and PU had disappeared from that of the spectrum of PU-dECM/PCL fibers.

#### 2.2.2. Immunofluorescent imaging for ECM protein detection on the PU-dECM/PCL fiber surface

To support the FT-IR analysis results, immunostaining was conducted as a qualitative assessment, to confirm the presence of dECM in the sheath of PU-dECM/PCL fibers. Fluorescent images were obtained for both PU-dECM/PCL and control PU-PCL nanofiber samples stained with five major proteins commonly found in cardiac dECM - collagen I, collagen III, elastin, fibronectin, and laminin. Fluorescence was observed only in the images of PU-dECM/PCL fibers (**Figure 4a-e**) whereas those of the control PU-PCL fibers appeared dull and dim (**Figure 4f-j**). All PU-dECM/PCL samples exhibited the expected fluorescence: collagen I - green (**Figure 4a**), elastin - green, (**Figure 4c**), collagen III - red (**Figure 4b**), laminin - red (**Figure 4e**), and fibronectin - blue (**Figure 4d**).

**Figure 4.**
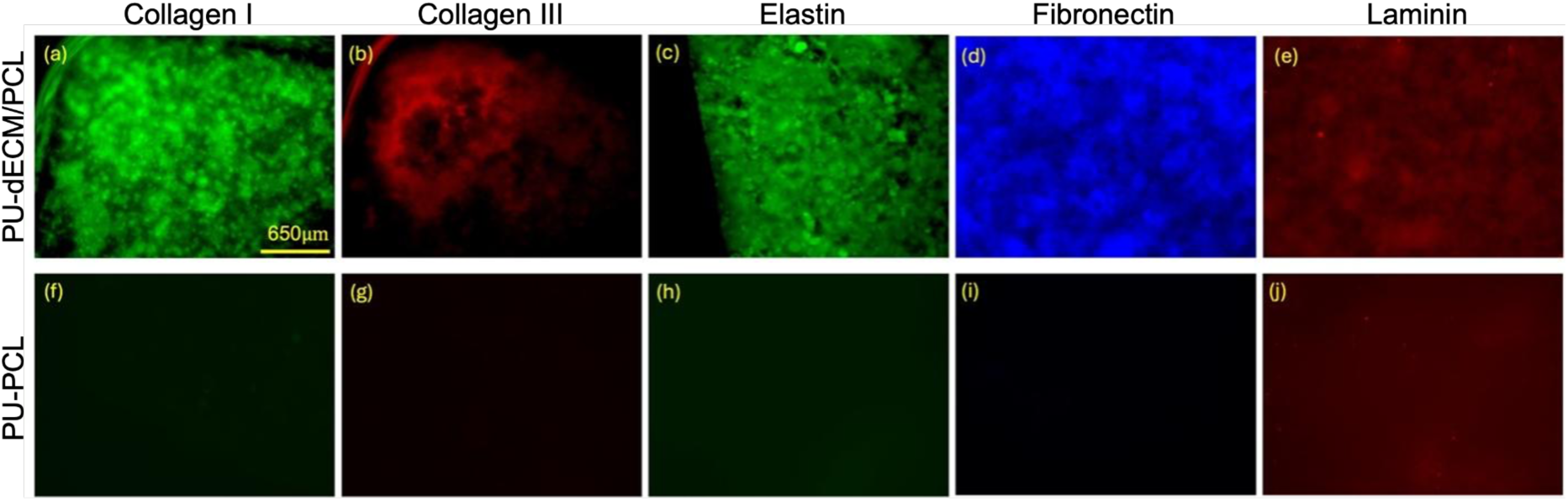
Confirmation of the presence of ECM proteins on the PU-dECM/PCL fiber surface. Immunofluorescent images of the coaxially electrospun PU-dECM/PCL and PU-PCL fiber specimens stained for collagen I (a) and (f), collagen III (b) and (g), elastin (c) and (h), fibronectin (d) and (i), laminin (e) and (j). Scale bar = 650μm

### 2.3. Characterization of the surface hydrophilicity of the three electrospun nanofiber types

The surface hydrophilicity of the fibers was evaluated by measuring the average contact angles with a 3 µl water droplet (**Table 3**, **Figure S2**). Both top (the surface on which fibers got deposited continuously during electrospinning) and bottom (the surface that touched the copper shim during electrospinning) surfaces of the fiber samples were evaluated.

**Table 3.** Calculated average contact angle values for PU-dECM/PCL, dECM/PCL, and PU-PCL nanofiber samples (mean ± standard error of the mean).

| Surface | Average contact angle ± SEM [°] | | |
| --- | --- | --- | --- |
|  | PU-dECM/PCL | dECM/PCL | PU-PCL |
| Top | 82.85 ± 1.5 | 79.92 ± 1.6 | 129.97 ± 0.41 |
| Bottom | 71.28 ± 1.8 | 72.2 ± 1.2 | 129.93 ± 0.38 |

It was noted that both surfaces of PU-dECM/PCL and dECM/PCL fiber samples produced contact angles less than 90°, whereas those of PU-PCL samples produced contact angles greater than 90° which were significantly higher (***p < 0.001). The contact angles of top surfaces of the samples with dECM (both monolith and core-sheath fibers) were not statistically significant (p = 0.189) and a similar result was observed with their bottom surfaces as well (p = 0.669) meaning their overall hydrophilicities were approximately similar. However, the difference between the contact angles produced by the top and bottom surfaces of these two sample types were statistically significant (***p < 0.001). In contrast, no such significant difference was noted in the two surfaces of PU-PCL samples (p = 0.934).

### 2.4. Characterization of the mechanical properties of the three electrospun nanofiber types

Mechanical properties of the three fiber types were evaluated by conducting tensile tests. Accordingly, respective stress-strain curves were obtained (**Figure 5**) and the average values of Young’s modulus, ultimate tensile strength (UTS), and % strain at UTS were calculated. These results are expressed as mean ± standard error of the mean in **Table 4**.

**Figure 5.**
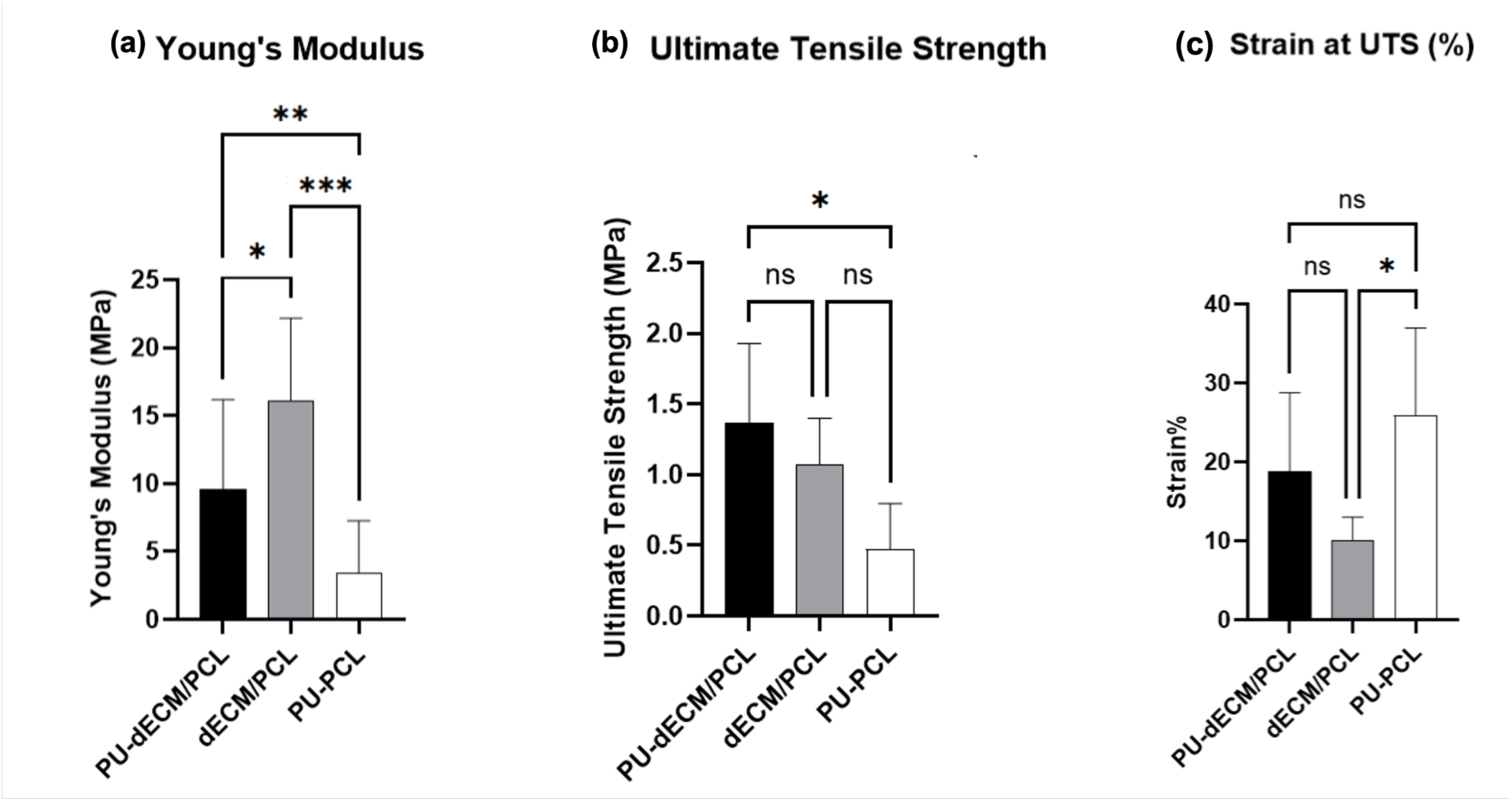
The comparison of the mechanical properties of PU-dECM/PCL, dECM/PCL, and PU-PCL fiber types. (a) Young’s modulus (b) Ultimate Tensile Stress (c) % strain at UTS. Significance Scale: Not Significant (ns): p > 0.05, Marginal Significance (*): 0.01 ≤ p ≤ 0.05, Moderate Significance (**): 0.001 ≤ p < 0.05, High Significance (***): 0.0001 ≤ p < 0.001.

**Table 4.** Mechanical properties of the PU-dECM/PCL, dECM/PCL, and PU-PCL fiber samples (mean ± standard error of the mean).

| Mechanical property | PU-dECM/PCL | dECM/PCL | PU-PCL |
| --- | --- | --- | --- |
| Young's modulus [MPa] | 9.57 ± 2.09 | 16.08 ± 1.93 | 2.15 ± 0.24 |
| Ultimate tensile strength [MPa] | 1.36 ± 0.25 | 1.07 ± 0.15 | 0.47 ± 0.14 |
| % strain at UTS | 18.86 ± 4.44 | 10.03 ± 1.34 | 25.87 ± 4.96 |

dECM/PCL exhibited the highest Young’s modulus, followed by PU-dECM/PCL and PU-PCL fibers. However, PU-dECM/PCL fibers had the highest UTS with dECM/PCL and PU-PCL coming next. In terms of the % strain at UTS, the values decreased in the order of PU-PCL, PU-dECM/PCL, and dECM/PCL. A significant increase in Young’s modulus was observed in the fibers with dECM compared to the purely synthetic PU-PCL fibers with core-sheath structure (**p = 0.007 with PU-dECM/PCL and ***p < 0.001 with dECM/PCL). However, core-sheath fibers containing dECM were significantly less stiff than the monolithic fibers with dECM (**p = 0.035).

### 2.5. Analysis of the enzymatic degradation behavior of the three electrospun nanofiber types

To approximately understand how the scaffold would degrade *in vivo*, an *in vitro* enzymatic degradation study was conducted. A mixture of collagenase – type I and lipase, was used to investigate the enzymatic degradation of collagen present in dECM and the two synthetic polymers: PCL and PU respectively.

Specimens of the three fiber types were immersed in the enzymatic solution as described in the methods section. Over four weeks, the mass and morphological changes were observed and presented as changes in their microstructure and mass (**Figures 6**). Notable surface cracks or flakes were observed in the SEM images of all fiber types, which are indicated by arrows in **Figure 6**. In samples containing dECM, those changes were observed from day 7 onwards (**Figure 6c, d, e, f, i, j, k, l**), while those weren’t visible in PU-PCL fibers until day 21 (**Figure 6q, r**).

**Figure 6.**
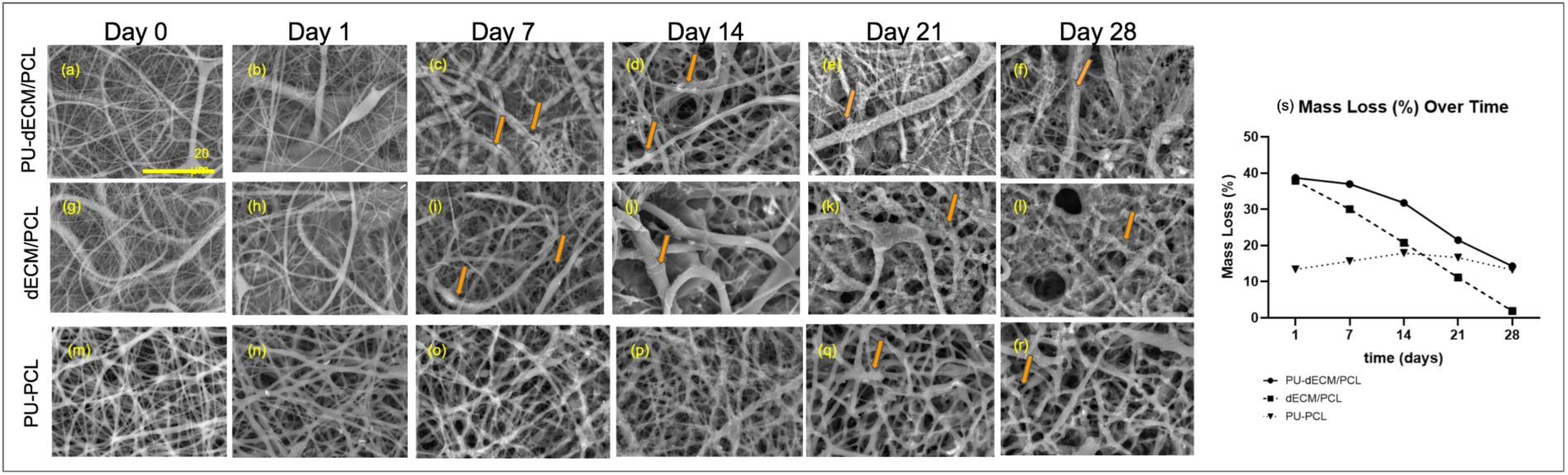
Qualitative and quantitative assessment of the enzymatic degradation of the three fiber types. SEM images of the (a – f) PU-dECM/PCL, (g – l) dECM/PCL, and (m – r) PU-PCL fiber specimens, obtained over the course of 28 days of the degradation study, at x2500 magnification, scale bar = 20 μm, % change in mass of PU-dECM/PCL, dECM/PCL, and (s) PU-PCL fibers over time in the presence of collagenase and lipase.

We observed the percentage mass loss over a four-week study (**Figure 6s**). The initial mass of each sample served as the reference point, with subsequent reductions indicating progressive degradation. PU-dECM/PCL fibers exhibited an initial mass loss of approximately 40%, followed by a steady decline, reaching around 20% after 28 days. Similarly, dECM/PCL fibers began with a comparable mass change (∼40%) but demonstrated a more rapid degradation, decreasing to nearly 0% by the end of the study. Both fiber types containing dECM showed significant mass reduction over time, with dECM/PCL fibers degrading at a faster rate. Among all tested samples, dECM/PCL fibers exhibited the most pronounced mass loss, whereas PU-PCL fibers displayed the least variation over the study period.

### 2.6. Cytotoxicity analysis of the three electrospun nanofibrous scaffolds in the presence of iPSC-CM

Given that the dECM was derived from the myocardial region, iPSC-CMs were employed to evaluate biocompatibility via LIVE/DEAD staining. This study aimed to assess the influence of native myocardial proteins on the retention and viability of cardiac cells. The heterogeneous cell population resulting from iPSC-CM differentiation offers a physiologically relevant model to investigate the role of native myocardial matrix components in CM adhesion and survival. Furthermore, this system enables the study of post-myocardial infarction (MI) remodeling, where native cardiomyocyte loss often promotes fibroblast proliferation—a process that can also be modeled using the iPSC-CM platform, as demonstrated here.

iPSC-CMs were generated using a previously established Wnt signaling modulation protocol^[30]^. On day 12 of differentiation, contractile iPSC-CMs were seeded onto three types of nanofibrous scaffolds, with TCPS serving as the positive control. Across a 7-day culture period, the TCPS group showed comparable cell viability to the dECM-containing scaffolds (**Figure 7**). However, the PU-PCL scaffold, which lacked dECM, exhibited a distinct decline in cell viability and altered morphology.

**Figure 7.**
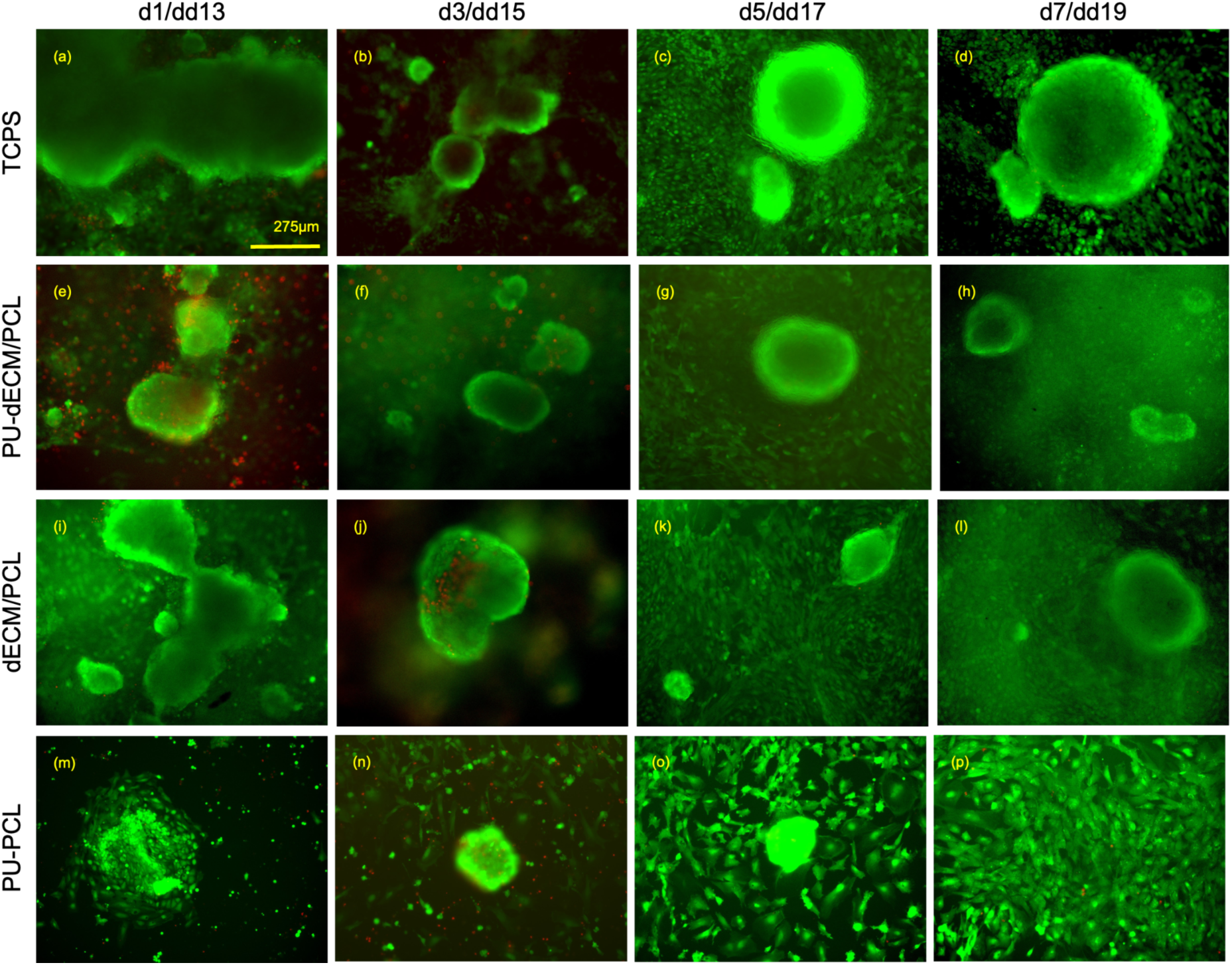
Live/Dead Staining of iPSC-CMs on the three nanofibrous scaffold types. (a-d) TCPS - cells retain cardiomyocyte morphology, consistent with native conditions. (e-h) PU-dECM/PCL scaffolds - cells exhibit a rounded, aggregated CM phenotype. (i-l) dECM/PCL scaffolds—cells maintain a rounded morphology characteristic of cardiomyocytes. (m-p) PU-PCL scaffolds - cells display an elongated, fibroblast-like morphology, indicating potential dedifferentiation. (Labels: *d* - days post iPSC-CM seeding on scaffolds. *dd* - days of differentiation after GSK3-inhibition, with dd0 as the starting point. Red fluorescence: dead cells. Green fluorescence: live cells, scale bar = 275µm.)

On day 1, cells on the PU-PCL scaffold initially appeared morphologically similar to other groups. By day 3, a reduction in cell density was evident, and by day 7, the majority of cells displayed a fibroblast-like, elongated morphology. This contrasted with the rounded, aggregated morphology typical of CMs, observed in both dECM-containing scaffolds and TCPS controls. The elongated phenotype on PU-PCL suggests potential dedifferentiation of iPSC-CMs, likely due to the absence of native ECM proteins in the scaffold. These findings highlight the importance of dECM in maintaining the structural and functional phenotype of iPSC-CMs in engineered scaffold environments.

### 2.7. ICC analysis of iPSC-CMs on the three electrospun nanofibrous scaffolds

Scaffolds seeded with iPSC-CMs were analyzed via ICC staining for major cardiac markers, such as alpha actinin (ACTN2) and troponin T (TNNT2). ACTN2 is an actin-binding compound that is present in the cytoskeleton of calcium-sensitive muscle cells, such as CMs. They have an essential function in stabilizing the contractile muscle apparatus (actin and myosin), connecting proteins in the cytoskeleton, which are used in the contractility of the cells. TNNT2, another marker in the cardiac cells, is a compound belonging to the troponin complex, which regulates the interaction between actin and myosin, which are responsible for the contractility of the CM population. When stained for ACTN2 and TNNT2, the scaffolds containing dECM components showed higher expression of CM markers, confirming that the dECM components help enhance the retention of CM markers when compared to the samples without dECM (**Figure 8**).

**Figure 8.**
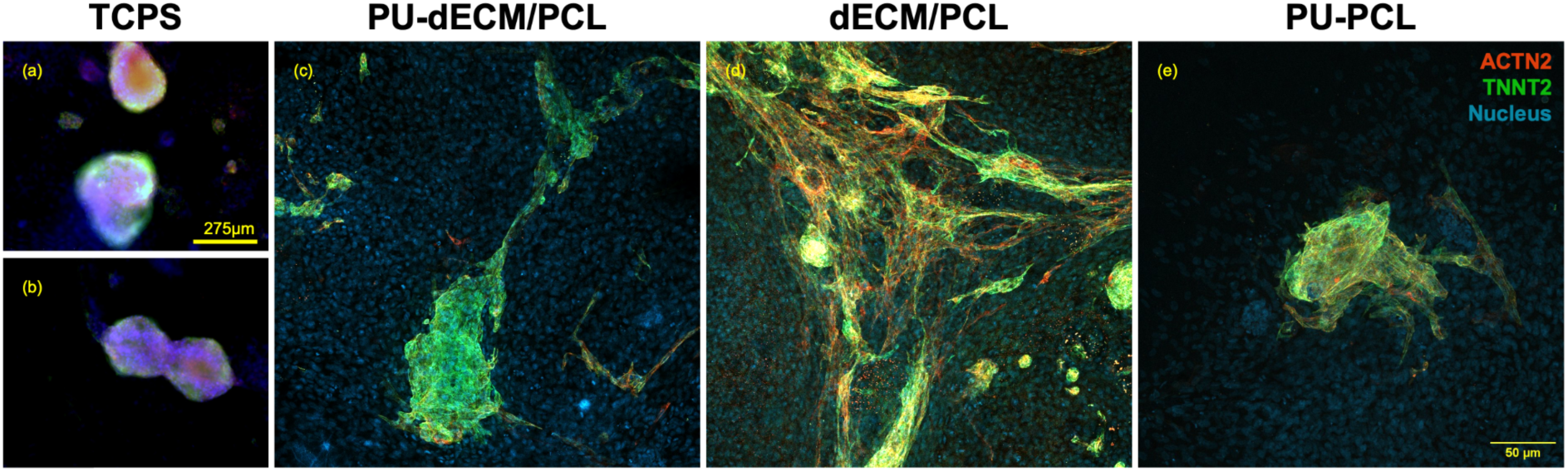
Immunostaining of iPSC-CM Specific Markers on the three nanofibrous scaffold types. Cardiac-specific markers were stained to assess iPSC-CM distribution and phenotype after 7 days of seeding on biomaterials. (a-b) Positive control on TCPS. (c) PU-dECM/PCL. (d) dECM/PCL. (e) PU-PCL. Among the tested materials, dECM/PCL scaffolds exhibited the most extensive cell spread, with strong expression of cardiac troponin T (green) and alpha-actinin (red), indicating enhanced cardiomyocyte retention and maturation.

The samples containing monolithic dECM fibers exhibited the most extensive distribution of cardiomyocyte-specific markers ACTN2 (red) and TNNT2 (green), suggesting that the dECM provides a favorable microenvironment for iPSC-CMs to form a continuous contractile network. Notably, ACTN2 expression was particularly pronounced in these samples, further supporting the role of dECM in promoting sarcomeric organization. In contrast, while the PU/PCL-dECM composite samples showed a broader spatial distribution of CM markers compared to the control, ACTN2 expression was more limited, likely due to the lower overall dECM content in the blend. Samples lacking dECM demonstrated the least marker expression and cellular spread, with iPSC-CMs confined to a localized region of the scaffold. This restricted distribution highlights the reduced biocompatibility of scaffolds without dECM and underscores the importance of dECM content in supporting iPSC-CM attachment and maturation.

### 2.8. Genetic Expression of iPSC-CM on the three electrospun nanofibrous scaffolds

RT-qPCR was conducted to evaluate the relative gene expression of CM markers across three scaffold groups, using iPSC-CMs cultured on TCPS as the control and GAPDH as the housekeeping gene. The expression levels of MYH7, TNNT2, and ACTN2 were assessed for each scaffold and analyzed using unpaired t-tests (**Figure 9**). On the PCL/PU scaffolds, MYH7 and TNNT2 expression levels were not significantly different from the control, while ACTN2 expression was significantly lower (****p < 0.0001). In contrast, the dECM-PCL/PU scaffolds showed significantly higher expression of TNNT2 relative to the control (***p < 0.001). Similarly, scaffolds composed of monolithic dECM fibers demonstrated significantly elevated expression of MYH7 (***p < 0.001), with ACTN2 (*p = 0.047) and TNNT2 (*p = 0.012) also significantly upregulated.

**Figure 9.**
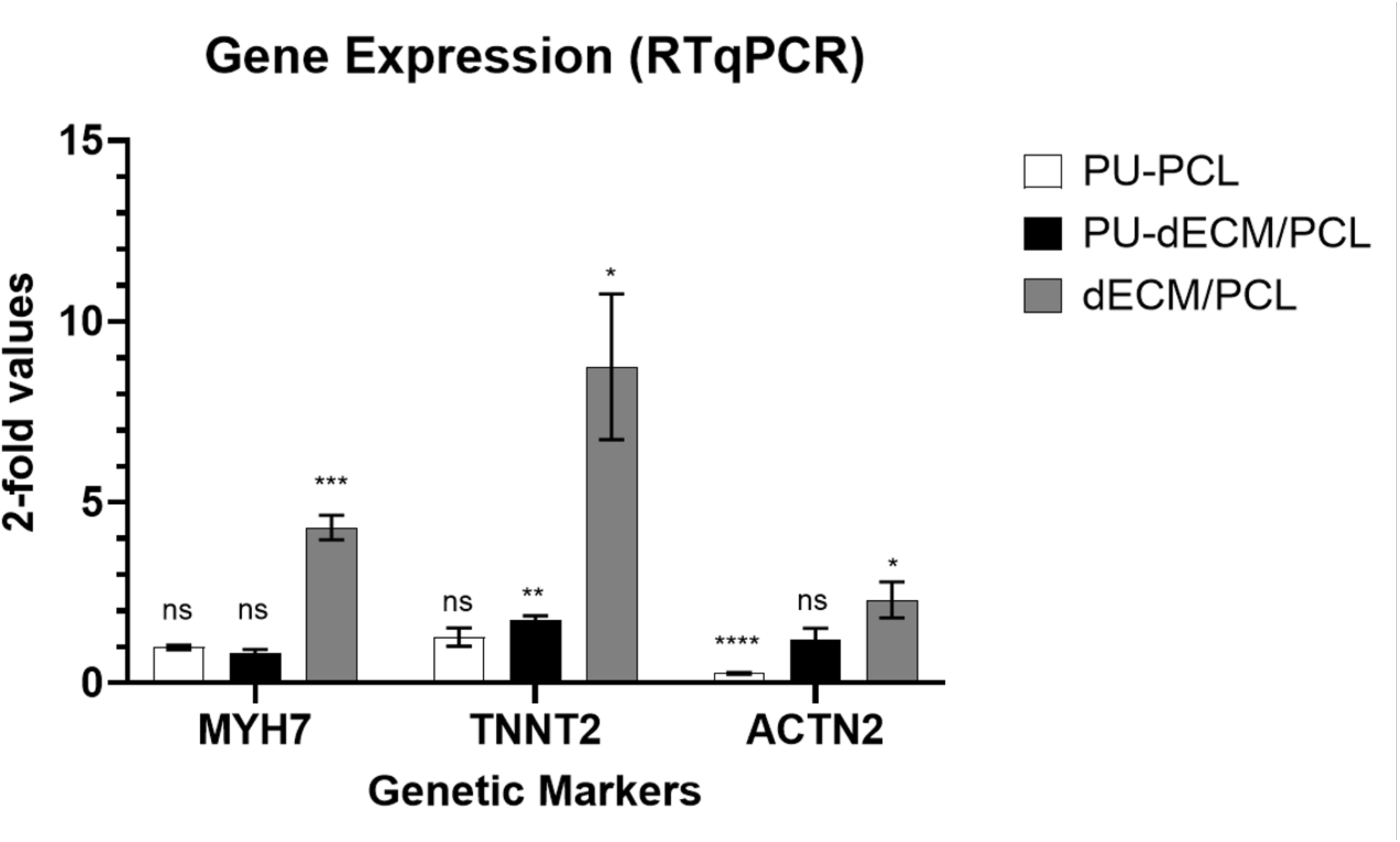
Genetic Expression of iPSC-CMs on Different Scaffolds. Expression levels of *Myosin Heavy Chain-7*, *Cardiac Troponin T*, and *Alpha Actinin* were analyzed in iPSC-CMs cultured on different scaffolds. iPSC-CMs on TCPS served as the control group, and a one-sample t-test was performed with a null hypothesis of 1. Significance Scale: Not Significant (ns): p > 0.05, Marginal Significance (*): 0.01 ≤ p ≤ 0.05, Moderate Significance (**): 0.001 ≤ p < 0.05, High Significance (***): 0.0001 ≤ p < 0.001, Very High Significance (****): p < 0.0001.

### 2.9. Functional Analysis of iPSC-CM on the three nanofibrous substrates

Calcium imaging revealed distinct functional differences in iPSC-derived cardiomyocytes (iPSC-CMs) across different substrate conditions (**Figure 10**). Cells cultured on tissue culture polystyrene (TCPS) exhibited the highest spontaneous beating rate, with an average BPM of 39.1 and a corresponding inter-beat interval (IBI) of 1.54 ± 0.06 s (*n* = 11). In contrast, cells on PU/PCL nanofibers demonstrated a significantly reduced BPM (10.9) and prolonged IBI (5.55 ± 0.47 s, *n* = 2), indicative of diminished automaticity.

**Figure 10.**
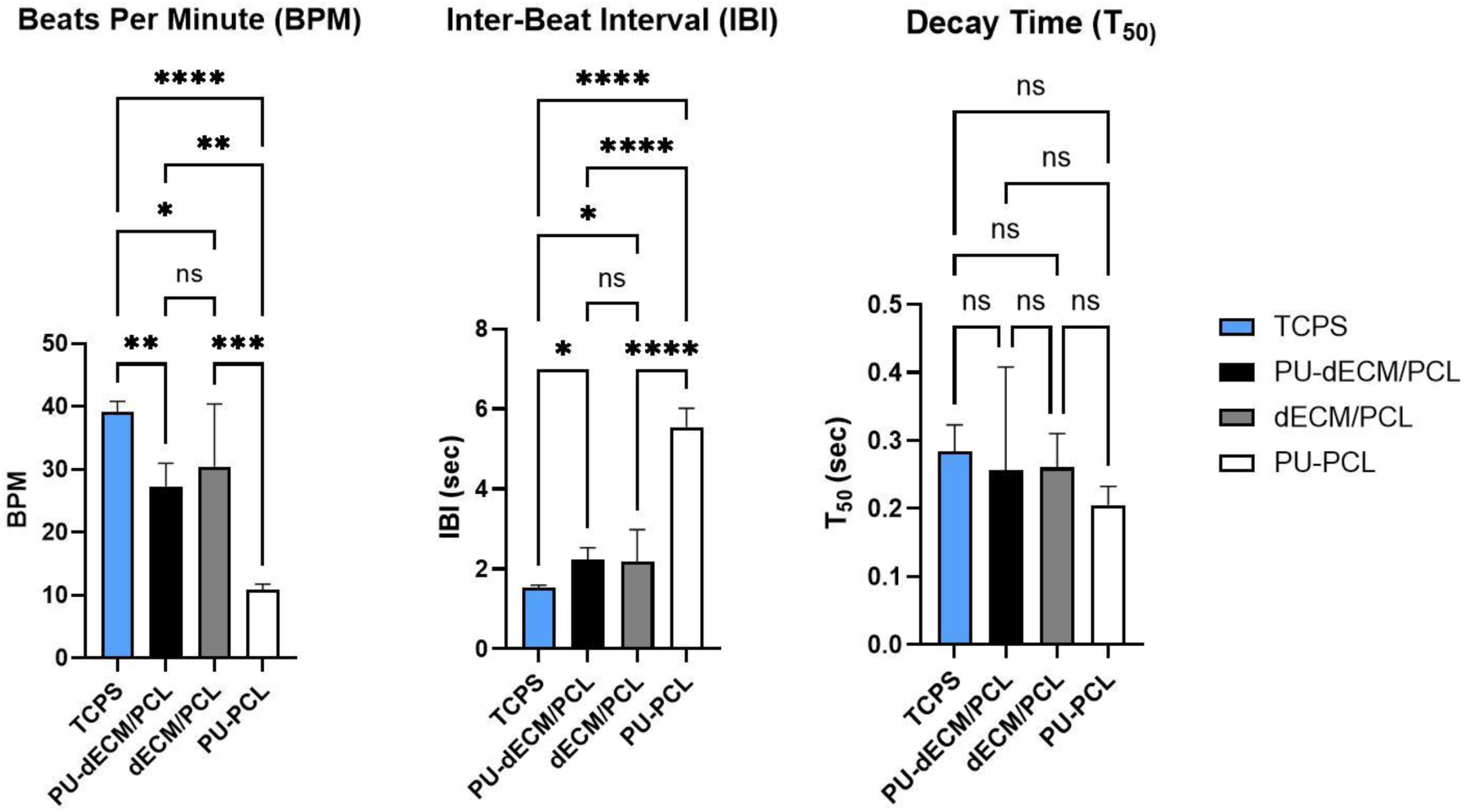
Functional Analysis of iPSC-CMs on samples with and without dECM. Contraction parameters like BPM, InterBeat Interval, and Time to 50% signal decay (T_50_) were analyzed in iPSC-CMs on monolithic dECM, PU-PCL/dECM, PU-PCL, and TCPS, and one way ANOVA was performed for significant differences. Significance Scale: Not Significant (ns): p > 0.05, Marginal Significance (*): 0.01 ≤ p ≤ 0.05, Moderate Significance (**): 0.001 ≤ p < 0.05, High Significance (***): 0.0001 ≤ p < 0.001, Very High Significance (****): p < 0.0001.

Introduction of decellularized extracellular matrix (dECM) has improved contractile function. PU/PCL-dECM samples showed intermediate values (BPM: 27.3; IBI: 2.23 ± 0.30 s, *n* = 5), while cells on dECM-coated substrates exhibited a BPM of 30.4 and IBI of 2.18 ± 0.80 s (*n* = 5) (**Figure 10**). These results suggest that dECM partially restores rhythmic beating in nanofiber systems and supports more regular activity compared to PU/PCL alone.

Decay kinetics of calcium transients, assessed as the time to 50% signal decay (T₅₀), were fastest in the TCPS group (0.28 ± 0.04 s, *n* = 7), followed by dECM (0.26 ± 0.05 s, *n* = 6), PU/PCL-dECM (0.26 ± 0.15 s, *n* = 5), and PU/PCL (0.21 ± 0.03 s, *n* = 2). While variability was greater in fiber-based groups, the overall decay times remained within a functional range, suggesting competent calcium reuptake across conditions.

Statistical analysis (one-way ANOVA) revealed significant differences in BPM and IBI among the four groups (*p* < 0.05), supporting the conclusion that substrate composition modulates both the frequency and kinetics of calcium handling in iPSC-CMs.

## 3. Discussion

Materials for the core and sheath regions of the fibers were selected to achieve the desired bioactive and physicochemical properties; a strong substrate with an elasticity comparable with the native heart tissue, and with the biological cues to promote iPSC-CM viability. To achieve a strong yet elastic fiber core matching the native heart tissue’s mechanical properties, medical-grade aliphatic poly(ether-urethane) (Selectophore™ PU), known for its use in biosensors and chemosensors was used^[31,32]^. Porcine cardiac-derived dECM, compositionally and physicochemically similar to human cardiac ECM, was selected for the sheath and blended with PCL to enhance electrospinnability and ensure the formation of continuous, uniform fibers.

First, the optimal electrospinning parameters for the dECM/PCL sheath solution were determined. Accordingly, 20 kV positive voltage, 10 kV negative voltage, 10 cm TDC, and a 1.5 ml/hr flow rate were selected. As stated by D. Han and A. J. Steckl, maintaining a stable core-sheath structure depends on the difference between the viscosities of the core and sheath solutions and the rates at which they are dispensed^[13]^. Most studies have reported this was possible when the viscosity and the flow rate of the sheath solution were higher than those of the core solution^[16]^. Accordingly, the core PU solution with a 2% w/v concentration was dispensed at a 0.75 ml/hr. Core-sheath interface tension is another factor influencing the formation of an unbroken core-sheath structure during coaxial electrospinning. To minimize its effect, HFIP was used as the solvent for both core and sheath solutions, following Nguyen et al. HFIP effectively solubilized dECM, and provided a solution with uniform charge distribution and enhanced electrospinnability, enabling continuous fiber formation^[16]^.

Reoptimization was needed for PU-PCL control samples, as the initial electrospinning parameters caused fiber fragmentation and distortion. The positive voltage was reduced to 8 kV and the negative voltage to 0 kV, while TCD, sheath flow rate, core flow rate, and solution concentrations remained unchanged. SEM imaging (**Figure S3**) confirmed improved morphology with lower voltage. Since the flow rate remained constant, the core-sheath structure was likely maintained despite the voltage adjustments.

TEM imaging confirmed the core-sheath structure in coaxially electrospun fibers, showing distinct core and sheath regions with clear color contrast (**Figure 1a**), unlike the control monolithic fibers (**Figure 1b**), which had a uniform shade throughout their cross-sections. This validated the successful formation of bicomponent fibers through coaxial electrospinning. Furthermore, the clear boundary between inner and outer regions of the PU-dECM/PCL fiber cross-sections indicated there was no miscibility between the core and sheath materials.

SEM analysis (**Figure 2**) showed that both core-sheath and monolithic fibers with dECM had similar diameter distributions with most fiber in the 290–440 nm range. The core-sheath fibers had a slightly larger average diameter than monolithic fibers, however, both were within the fiber diameter range of native collagen fibers (20–500 nm) and other reported electrospun dECM fibers^[4,33]^.The increased diameter of PU-dECM/PCL fibers was likely due to the firm PU core supporting the dECM/PCL sheath.

On average, a reduction of ∼30% in fiber diameter was observed in the coaxially electrospun fibers with dECM as opposed to those without. However, since the two fiber types were fabricated under different voltages, a key parameter affecting fiber size, this reduction cannot be solely attributed to the incorporation of dECM. A 12 kV voltage difference could significantly impact fiber dimensions, making it difficult to isolate the effect of dECM addition on fiber diameter although the other processing conditions remained unchanged.

Randomly arranged electrospun nanofibers create interconnected pore networks that facilitate cell infiltration, proliferation, gaseous and nutrient diffusion, and the removal of excretory matter of cellular functions, which are key for TE scaffolds^[6,7,9,10,18,34,35]^. Accordingly, the average pore size of the scaffold should be sufficiently larger than the differentiated cells to enable their migration and integration. While porosity (percentage of void space) is often reported in literature (50–90%), our study focuses on pore size, which directly impacts scaffold architecture and cellular behavior.

Both fiber types containing dECM had comparable average pore sizes with most pores belonging to the 30–125 μm² range. Significantly smaller pores in PU-PCL could likely be due to differences in fabrication voltages, which possibly increased the fiber diameter and reduced the pore size, leading to a denser structure. While adult cardiomyocytes (17–25 μm in diameter, 60–140 μm in length)^[36]^ are well-aligned, iPSC-CMs are smaller and more misaligned (5–10 μm diameter)^[37]^. Given our observed pore sizes, all three scaffold types provide sufficient cell infiltration pathways, supporting iPSC-CM integration and function.

ATR-FTIR analysis was conducted to confirm the presence of the dECM/PCL blend on the surface of PU-dECM/PCL nanofibers (**Figure 3**). The presence of dECM on the sheath was crucial in achieving a cell-preferred substrate with excellent biocompatibility. FT-IR spectra of dECM powder, PCL, and PU pellets were compared with that of coaxially electrospun PU-dECM/PCL nanofibers. It was revealed that most PCL peaks remained unchanged, confirming its presence on the fiber surface and indicating that electrospinning and dissolution in HFIP did not alter PCL’s structure. Two additional peaks at 1653 cm^−1^ and 1540 cm^−1^, corresponding to amide C=O stretching and N-H bending, were observed in PU-dECM/PCL fibers, but due to their presence in both PU and dECM spectra ^[26,29]^ it was difficult to identify the source of those peaks on the PU-dECM/PCL spectrum. The disappearance of the peak at 3302 cm⁻¹ for N-H stretching suggested possible chemical changes during solution preparation or electrospinning^[26,28,29]^.

Gao et al. reported using immunofluorescent staining to confirm the presence of dECM on the surface of the fibers they produced by electrospinning a solution of dECM and PCL^[38]^. Similarly, to further confirm dECM presence on the core-sheath fiber surface, immunofluorescent staining was performed for five key ECM proteins: collagen I, collagen III, elastin, fibronectin, and laminin (**Figure 4**). Decellularization ideally preserves ECM structural proteins while removing immunogenic components^[18]^. Prior studies identified collagen I (85%) and III (11%) as dominant cardiac ECM components, along with elastin, fibronectin, and laminin, which support structural integrity and intercellular signaling^[39–41]^. Immunostaining results (**Figure 4**) showed positive fluorescence in PU-dECM/PCL fibers but not in PU-PCL controls, confirming the presence of dECM in the sheath. Combined with FT-IR data, this validated the successful formation of core-sheath fibers with PU in the core and a dECM/PCL sheath.

Hydrophilicity is a key requirement for a TE scaffold, as it directly affects cell attachment and proliferation^[42]^. Hydrophilic scaffolds enhance adhesion and growth by allowing better diffusion of cell suspensions and providing focal adhesion points^[43,44]^. Contact angle measurements indicate surface wettability, with surfaces producing angles below 90° classified as hydrophilic and those ≥ 90° as hydrophobic^[45]^. To assess fiber surface wettability, contact angles were measured using a goniometer, and average values were calculated for both the top and bottom surfaces.

Both PU-dECM/PCL and dECM/PCL sample types produced contact angles below 90°, indicating hydrophilic surfaces. Additionally, this confirmed that the surface wettability was not affected by the core-sheath structure. A notable difference existed between the contact angles of the top (82.85 ± 1.5°) and bottom (71.28 ± 1.8°) surfaces of PU-dECM/PCL fiber samples with the former being higher and this observation was consistent with the results of the dECM/PCL fibers (top – 79.92 ± 1.6°, bottom - 72.2 ± 1.2°) but no notable difference was observed in PU-PCL samples (top – 129.97 ± 0.41°, bottom – 129.93 ± 0.38°). This variation may have resulted from the smooth texture created on the bottom surface of the sample due to its constant contact with the copper shim, which likely flattened any protrusions or micro-textures on the surface, resulting in a more even surface. The significant difference between samples with and without dECM confirmed that dECM incorporation significantly improved fiber hydrophilicity, consistent with previous studies^[38,46,47]^.

Mechanical strength is essential for TE scaffolds to withstand different in vivo forces and cyclic loads. Moreover, it is reported that the stiffness or the elasticity of a scaffold directly influences cell migration and differentiation^[48]^. Ideally, the scaffold’s Young’s modulus should match that of native tissue to prevent functional mismatches. The Young’s moduli of the fibers with dECM irrespective of the fiber structure were higher than that of those without dECM. This increase could be attributed to ventricular dECM used, which is typically stiff to support the ejection of blood. A similar trend was reported by Feng et al., where fibers with a blend of PCL and decellularized ECM from rat auricular cartilage exhibited a fivefold increase in stiffness compared to pure PCL fibers^[46]^.

The reported elastic modulus of native myocardium varies widely. Healthy myocardium typically ranges from 3–100 kPa,^[39]^ with specific values reported as 35 kPa, below 50 kPa, and between 10–500 kPa depending on the cardiac cycle^[49,50,51]^. Given this, CTE scaffolds should ideally fall within the kilopascal range. However, all three tested fiber types had moduli in the megapascal range, exceeding native heart stiffness but aligning with FDA-approved cardiovascular implants such as vascular grafts (17.4 MPa), cardiac sutures (GPa range), and synthetic heart valves (10–100 MPa)^[52]^. As these materials are clinically successful, the stiffness discrepancy is unlikely to cause adverse effects when in use.

PU-PCL fibers exhibited the highest % strain at UTS, followed by PU-dECM/PCL and dECM/PCL, indicating greater ductility in fibers with the PU-core fibers (**Figure 5**). The modulus of the core-sheath fibers with dECM was 1.7 times lower than that of monolithic dECM fibers. Accordingly, the introduction of the PU core reduced the stiffness of the dECM/PCL fibers approximately by 40%. This confirms the potential of coaxial electrospinning in modulating the mechanical properties of nanofibers, balancing strength and flexibility for potential TE applications.

Degradability is a key requirement for TE scaffolds and is influenced by functional group stability, molecular weight, and environmental factors such as temperature, humidity, and pH^[19]^. Ideally, the scaffold’s degradation rate should match the tissue regeneration rate. To approximate in vivo degradation, an in vitro enzymatic degradation study was performed using collagenase type I and lipase to target collagen in dECM and synthetic polymers (PCL and PU), respectively (**Figure 6**).

Collagenase type I (MMP-I) effectively degrades collagen types I, III, and VII through proteolysis^[24]^. Alberti and Xu reported using 1 mg/ml collagenase concentration to analyze the degradation behavior of a TE scaffold composed of tendon-derived collagen fibrils^[24,52]^. Suh et al. reduced that concentration to 0.25 mg/ml due to the significant size difference between their CNT - electrospun PCL fibers and the tendon-derived collagen fibrils. The core-sheath fibers fabricated in our study were even smaller (approximately by 2 times) than Suh et al.’s. However, the same collagenase concentration was used in this study as it remains physiologically relevant, given that MMP levels increase post-myocardial infarction^[53]^. The lipase concentration (0.0025 mg/ml) was also chosen based on Suh et al.’s study, where a higher enzyme concentration was avoided to study degradation rather than accelerate it^[52]^.

SEM images revealed progressive fiber degradation, confirming enzymatic action. The observed increase in mass over time in PU-dECM/PCL and dECM/PCL fibers is attributed to swelling resulting from dECM-enhanced water absorption and liquid retention due to porosity of the samples (**Figure 6s**). Fiber swelling was observed in the SEM images of these two fiber types taken at each time point (**Figure 6a-l**). In contrast, PU-PCL fibers didn’t exhibit swelling, however, a mass increase was noted from day 7 to 14. This too could be due to the retention of liquid in the porous structure of the sample despite their inherent hydrophobicity. The decrease in the mass of PU-PCL fibers after day 14 indicated gradual material breakdown.

The reduction in the % mass change over time in the samples with dECM indicated their degradation, which caused both material loss and the loss of mechanical integrity of the interconnected porous network, reducing their ability to absorb and retain liquid. These trends were consistent with SEM observations, confirming that dECM incorporation enhanced degradation in PU-dECM/PCL and dECM/PCL fibers. The PU core in PU-dECM/PCL fibers likely preserved porous network integrity better than monolithic dECM/PCL fibers.

Since post-MI tissue regeneration typically takes 6–8 weeks, a scaffold should degrade within this timeframe to prevent inflammation^[21]^. By week 4, PU-dECM/PCL fibers exhibited ongoing degradation, suggesting they would continue degrading post-implantation. This study confirms that PU-dECM/PCL fibers undergo enzymatic degradation, supporting their potential as a scaffold for CTE applications.

The cell response was evaluated using differentiated iPSC-derived cardiomyocytes (iPSC-CMs) at day 9-12 of differentiation. Cells were initially differentiated on TCPS coated with laminin until spontaneous contractile activity was observed, after which they were seeded onto sterilized scaffold samples: dECM/PCL, PU-PCL, and PU-dECM/PCL. Within 24 hours, clusters of iPSC-CMs began to disaggregate into smaller clusters and single cells with elongated morphology, most prominently on PU-PCL scaffolds. By day 7 post-seeding, nearly all cells had assumed an elongated, fibroblast-like morphology. In contrast, substrates containing dECM retained more cell clusters and displayed visible contractile activity.

Immunostaining confirmed that samples retaining cluster morphology also exhibited elevated expression of cardiomyocyte-specific markers troponin T (TNNT2) and alpha-actinin (ACTN2), indicating improved retention of cardiac identity (**Figure 8**). We propose that the absence of dECM limits the availability of tissue-specific matrix cues, leading to dedifferentiation of iPSC-CMs toward their fibroblast origin, driven by epigenetic memory. This behavior aligns with literature showing that iPSCs retain somatic origin-specific gene expression patterns, which can influence lineage specification. For example, fibroblast- and cardiac progenitor-derived iPSCs differ in the timing of cardiac marker expression such as NKX2-5, although this distinction diminishes with continued passaging ^[54,55]^.

The dedifferentiation of cells is attributed to various factors including improper culture techniques or immature phenotype of the iPSC-CMs^[55]^. For the current study, we used early iPSC-CMs from days 9-12, a period when cells are in a plastic state and are susceptible to dedifferentiation under unsustainable conditions and the absence of native proteins induced additional stress, leading to dedifferentiation. Validation through staining and PCR revealed higher cardiac markers (TNNT2, ACTN2, and MYH7) in samples with dECM (PCL-dECM/PU and monolithic dECM/PCL) compared to samples without dECM (PU-PCL) (**Figures 8 and 9**).

To further evaluate cardiac functionality, calcium imaging was performed across all groups (**Figure 10**). Cells cultured on TCPS showed the highest spontaneous activity, with an average BPM of 39.1 and interbeat interval (IBI) of 1.54 ± 0.06 s. In contrast, PU-PCL scaffolds exhibited significantly reduced activity (BPM: 10.9; IBI: 5.55 ± 0.47 s), indicating poor contractile function and potential dedifferentiation. Scaffolds incorporating dECM demonstrated partial restoration of functional activity. Both PU/PCL-dECM and dECM-coated samples showed intermediate BPM values (27.3 and 30.4, respectively) and shorter IBIs (2.23 ± 0.30 s and 2.18 ± 0.80 s), suggesting improved cell retention and preservation of the cardiomyocyte phenotype. Calcium decay kinetics (T₅₀) were also assessed, with the shortest decay observed in TCPS (0.28 ± 0.04 s), and comparable decay times in the dECM groups (0.26 ± 0.05 s for dECM and 0.26 ± 0.15 s for PU/PCL-dECM), indicating competent calcium handling despite the scaffold transition.

It is important to note that the iPSC-CMs used in this study were initially differentiated on TCPS and subsequently dissociated before seeding onto scaffolds. This process, which typically involves enzymatic reagents such as ReLeSR or TrypLE, disrupts cell-cell junctions, particularly gap junctions and desmosomes, that are essential for coordinated cardiomyocyte function. Disruption of proteins like connexin 43 during dissociation has been shown to impair functionality, with studies demonstrating that restoring gap junction integrity leads to improved iPSC-CM performance ^[56,57]^. In the context of this study, the observed recovery of function in dECM-containing scaffolds may be attributed to the supportive role of native ECM proteins in promoting the reformation of cell–cell contacts and stabilizing cardiomyocyte function. Future work will include comparison of iPSC-CMs dissociated and replated on TCPS as a control as well as analyzing the CM function and presence of CM-specific markers for a longer time point to allow for additional time for cells to reestablish cell-cell and cell-matrix after dissociation.

In this study, the use of early-stage iPSC-CMs (day 9-12) may have increased susceptibility to dedifferentiation due to their plasticity and immature phenotype. When cultured in the absence of native matrix proteins, these cells experienced additional environmental stress, contributing to their reversion toward a non-cardiac lineage. Validation through immunostaining, functional calcium imaging, and PCR confirmed higher expression of cardiac markers (TNNT2, ACTN2, and MYH7) in dECM-containing samples (PU/PCL-dECM and monolithic dECM/PCL) compared to samples without dECM (PU-PCL), further supporting the conclusion that ECM composition is a critical modulator of iPSC-CM identity and function. This was further confirmed in the functional testing of the cells, where cells showed comparable beats per minute (BPM), interbeat interval (IBI), and decay time (T_50_) on PU-dECM/PCL and dECM/PCL, however the samples containing only synthetic polymer (PU-PCL) showed irregular interbeat interval, further validating the claim that dECM proteins help with restoring the functionality of iPSC-CMs by providing the native microenvironment to the cells compared to TCPS and synthetic polymers.

Although preliminary, the results provide confidence that the scaffold with core-sheath structure provides the structural and biochemical support to the cells when native proteins are incorporated into the structure. However, the content of dECM remains a highlight of this study, as more native dECM content helps with the structural spread of cells, providing higher signs of biocompatibility, whereas the functionality of cells remain unhindered by the dECM content of the material.

Tensile test results confirmed the improvement in mechanical properties due to the inclusion of a PU core into the dECM/PCL blended fibers, especially in terms of reducing their stiffness and also in achieving long-term stability in culture media. Biocompatibility results indicated that both fiber types containing dECM had comparable biological performance. It should be noted that the amount of dECM in the core-sheath fibers is less compared to that in the monolithic fibers. Therefore, we believe that increasing the dECM content in the sheath of the coaxially electrospun fibers could further enhance cellular response while maintaining the improved mechanical properties provided by the core-sheath structure.

## 4. Conclusion

This study focused on using coaxially electrospun core-sheath nanofibers for cardiac tissue engineering applications, where decellularized myocardial extracellular matrix (dECM) was used in conjunction with polycaprolactone (PCL) and polyurethane (PU). This natural-synthetic material combination was used to enhance the physical properties and biological performance of the fibrous scaffold thereby improving its mimicry of the native cardiac microenvironment. Accordingly, a blend of dECM and PCL was used in the sheath to provide the biological cues necessary for cell growth and survival whereas the PU core was used to provide adequate elasticity for the scaffold.

With optimized electrospinning conditions, we obtained nanofibers with sizes comparable to collagen fibers in the natural ECM. TEM image analysis confirmed the formation of fibers with the core-sheath geometry and FT-IR analysis together with immunostaining confirmed the relative location of the three constituents - dECM and PCL on the sheath and PU in the core. Interestingly, we observed the improvement in hydrophilicity and the rate of degradation due to the presence of dECM. Tensile testing results confirmed that the core-sheath fiber structure helped reduce the stiffness of the blended monolith fibers, which demonstrated the ability of coaxial electrospinning to modulate the mechanical properties of fibers by deploying the suitable materials. When seeded with iPSC-CM, the material showed comparable biocompatibility results with the monolithic dECM-based fibers (dECM/PCL). When compared to the positive control of iPSC-CM on TCPS, there was lesser dedifferentiation of the iPSC-CM, providing evidence that the coaxial nature of the fibers did not compromise the biocompatible nature of dECM.

There are multiple avenues through which this study could be continued. Since these fibers show excellent biocompatibility with iPSC-derived cardiomyocytes, they present a promising platform for the differentiation of iPSCs in vitro. Therefore, their influence on iPSC differentiation into cardiomyocytes could be investigated. In addition to that, it would be interesting to investigate how the scaffolds with different dECM concentrations and fiber morphologies (size, shape, arrangement) affect cellular behavior. Moreover, since this study has confirmed the applicability of coaxial electrospinning for creating dECM-based core-sheath fibers, the same fabrication method could be tried for making dECM core-sheath fibers containing other synthetic and/or natural material combinations and investigate their structure-property-performance relationship.

It is also possible to change the material composition to achieve different functional properties in the fibers. For instance, blending dECM with a conducting polymer with proven electrospinnability, such as polypyrole to form the sheath, conductivity could be induced on the fiber surface. Such a conductive bioactive scaffold could potentially enhance the biological performance of a scaffold for cardiac tissue engineering due to the inherent conductivity displayed by the native cardiac tissues to facilitate the contractile function. Furthermore, such multi-functional fibrous scaffolds could serve as a single platform to study how external stimulations, either one or a combination, could influence cell behavior by using them in appropriate bioreactors. Additionally, the core-sheath fiber configuration could be used for creating drug-eluting scaffolds with both regenerative and therapeutic properties.

## 5. Experimental Section/Methods

### 5.1. Decellularization of ECM extracted from porcine heart

Porcine hearts were acquired from Nahunta Pork Center (Raleigh, NC) and were stored at −80 °C. Prior to decellularization, the hearts were cut into small pieces of about 125mm^3^. The samples were washed with tap water until the water ran clear. Subsequently, the samples went through a series of washes: deionized water (5 minutes), 2X DPBS (15 minutes), 0.02% Trypsin EDTA in 1X DPBS (120 minutes), deionized water (5 minutes), 2X DPBS (15 minutes), 3% Tween-20 in deionized water (120 minutes), deionized water (5 minutes), 2X DPBS (15 minutes), 4% w/v sodium deoxycholate in deionized water (120 minutes), deionized water (5 minutes), 2X DPBS (15 minutes), 0.1% v/v peracetic acid in 4% v/v ethanol in deionized water (60 minutes), 1X DPBS (5 minutes), deionized water (5 minutes), 1X DPBS (5 minutes), DNase-I (20 minutes). All washing was done at room temperature except for DNase-I, which was incubated at 37 °C. The samples were then lyophilized for 72 hours, followed by grinding into powder form.

### 5.2. Preparation of solutions for electrospinning the three nanofiber types

To obtain the sheath solution to fabricate PU-PCL/dECM samples, first, two separate solutions of 14% (w/v) PCL and 6% (w/v) dECM were prepared by dissolving 700mg of PCL (Mw 150k kDa) pellets in 5ml of HFIP and 300mg of dECM in another 5ml of HFIP respectively. The two solutions were stirred overnight on a magnetic stir plate at 600 rpm. Then these two solutions were mixed in a 1:1 ratio to make a polymer solution with a total concentration of 10% (w/v). The resultant solution was stirred for an additional day before being used for electrospinning. For the core, a 2% (w/v) Polyurethane solution was prepared by dissolving 200mg of PU in 10 ml of HFIP. The solution was stirred overnight on a magnetic stirrer at 600 rpm before being used for electrospinning.

To fabricate the PU-PCL control samples, the same core solution was used whereas a 10% (w/v) PCL solution was prepared as the sheath solution by dissolving 1g of PCL in 10 ml of HFIP. This solution was stirred overnight on a magnetic stirrer at 600 rpm before being used for electrospinning. To fabricate the PCL/dECM control samples, the 10% (w/v) sheath solution used for PU-PCL/dECM samples was used. **Table 5** below shows the polymers and their concentrations in the solutions used for synthesizing the respective samples.

**Table 5.**
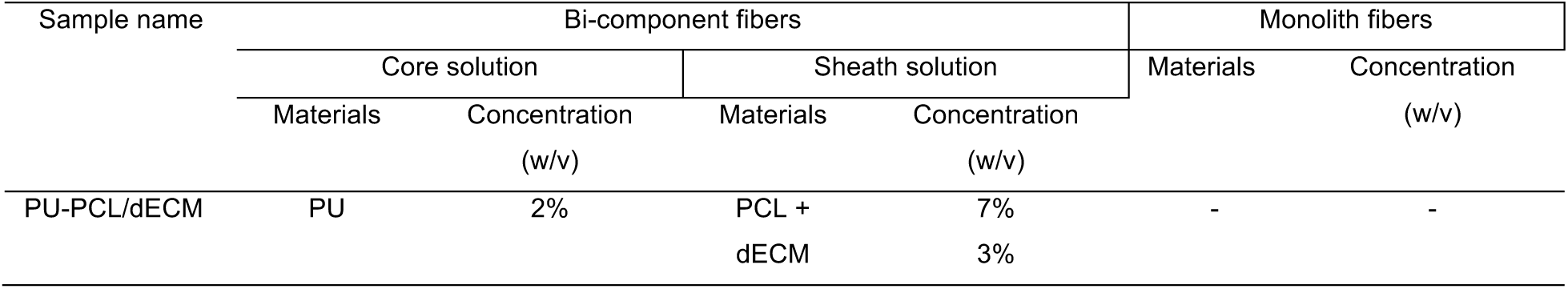
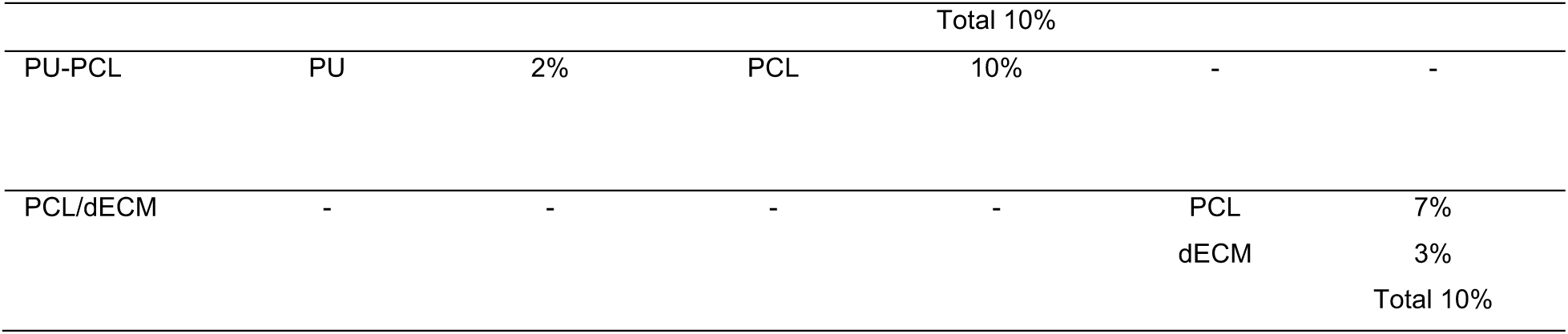
Ingredients and concentrations of the solutions used to fabricate PU-dECM/PCL, dECM/PCL, and PU-PCL fiber samples.

### 5.3. Electrospinning of the three nanofiber types

The coaxial electrospinning setup (**Figure 11a**) was used to fabricate both PU-PCL/dECM samples and PU-PCL control samples.

**Figure 11.**
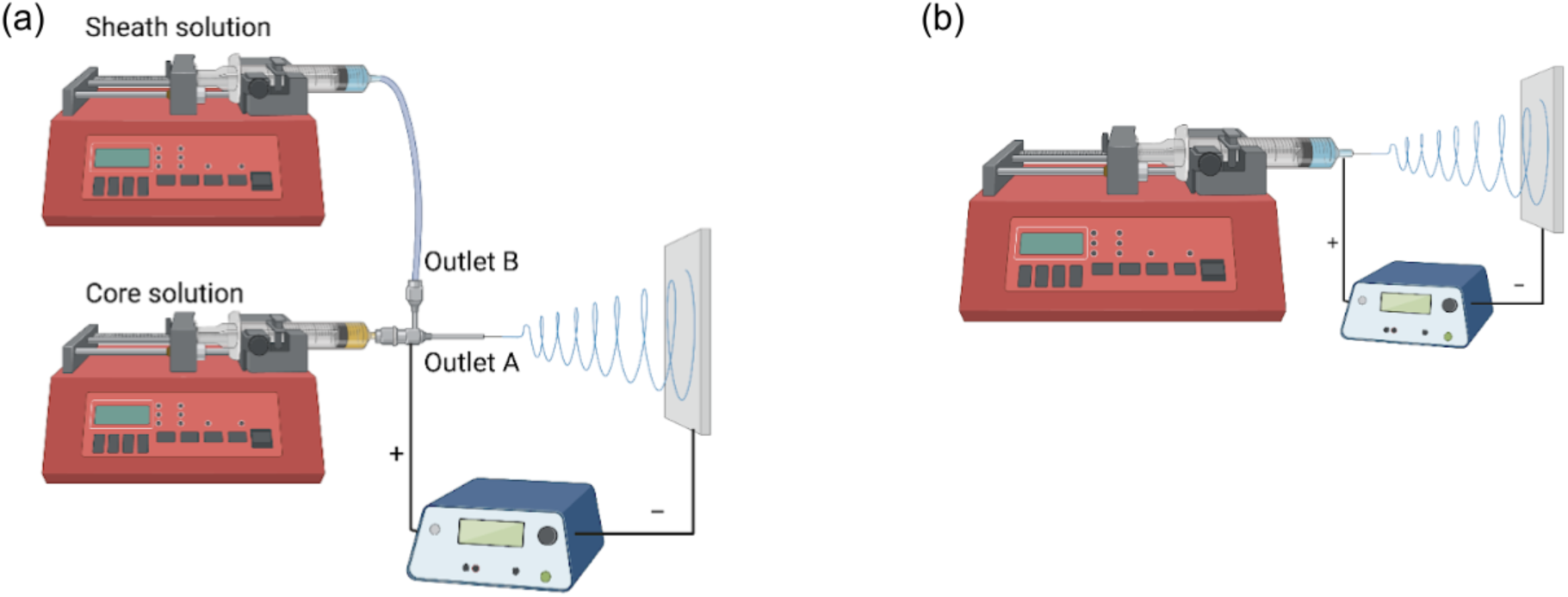
Schematics of electrospinning set-ups. (a) coaxial electrospinning and (b) conventional monoaxial electrospinning

First, 2ml of the core solution was loaded to a syringe, to which then the coaxial needle was fixed. It was then mounted onto a syringe pump, in front of which the copper collector was placed. The distance between the needle tip (outlet A as shown in **Figure 11a**) and the copper collector was maintained at 10cm. Next, 2ml of the sheath solution was loaded to another syringe. This was connected to the coaxial needle (to outlet B as shown in **Figure 11a**) via a 12 cm long narrow tube with an inner diameter of 1 mm. Electrospinning was initiated by supplying a positive voltage of 20 kV to the needle and a negative voltage of 10kV to the collector as the core solution was extruded at a rate of 0.75 ml/ hr and the sheath solution at 1.5 ml/hr. When electrospinning PU-PCL fibers, the positive and negative voltages were lowered to 8kV and 0kV respectively as the previous voltage produced a higher number of irregular, discontinuous fiber fragments. All the other process parameters were maintained the same without any change.

The conventional electrospinning setup (**Figure 11b**) was used to fabricate the PCL/dECM samples. The voltage and the needle-to-collector distance were the same as in coaxial electrospinning whereas the flow rate was maintained at 1.5 ml/hr.

The electrospun fibers were collected as mats on the stationery copper shim and then deposited in a desiccator to let any residual solvent evaporate and dry the samples up for subsequent characterizations. The room temperature and the relative humidity were maintained at 22 – 25 ^0^C and 27 – 35% during electrospinning. Fibrous meshes with thicknesses of 0.4 ± 0.03 mm, 0.42 ± 0.04 mm, and 0.38 ± 0.029 mm were obtained for PU-dECM/PCL, dECM/PCL, and PU-PCL sample types, respectively.

### 5.4. Characterization of the structure and morphology of the three electrospun nanofiber types

#### TEM image analysis

TEM image analysis was done to confirm the presence of core-sheath geometry in the coaxially electrospun fibers. Cross-sectional images of the fibers in each sample type were obtained using a Hitachi HT7800 Transmission Electron Microscope (Japan). For cross-sectional imaging, 1 cm^2^ samples were prepared by resin embedding followed by sectioning with an ultra-microtome (Leica ultra-microtome EM UC7), Germany) and staining with osmium vapor overnight for better contrast. The samples of PU-PCL/dECM, and PCL/dECM fibers were observed under a 20,000x magnification. The presence of the core-sheath arrangement was confirmed by the color contrast observed in the inner and outer regions in the coaxially electrospun fibers, which was not apparent in the conventionally electrospun monolith fibers.

#### SEM image analysis

SEM images of the PU-PCL/dECM, PU-PCL, and PCL/dECM fibers were analyzed to study the morphological differences between them. 5 specimens of 1 x 1 cm^2^ were cut from each sample type and coated with a 10nm layer of gold and palladium using Emietch sputter coater to induce conductivity prior to imaging. Next, the specimens were observed using a scanning electron microscope (Hitachi TM4000) with 15kV accelerating voltage and BSE (backscattered electron) mode, under different magnifications.

6 images per specimen were obtained with 5000x magnification, and 10 measurements were taken from each image for fiber diameter (a total of 300 measurements) using Image J software (NIH). 4 images per specimen were obtained with 2500x magnification, and 5 measurements were taken from each image for pore size (a total of 100 measurements) using the same software. Those measurements were used to calculate the average values for fiber diameter and pore size for each sample type, as well as to create their respective distribution curves.

### 5.5. Analysis of the chemical composition of the three electrospun nanofiber types

#### FT-IR analysis to identify functional groups

FT-IR analysis was done using Thermo Fisher model iS10 with OMNI ATR sampler with Germanium (Ge) crystal to study the chemical composition of the core-sheath fibers and the relative location of each constituent. The spectra for individual constituent materials (PU pellets, PCL pellets, and dECM powder) and 1cm x 1cm PU-PCL/dECM fiber samples (n=3) were obtained between the wavelength range from 4000 – 400 cm^−1^ using the ATR mode. The functional groups indicated in the spectra for the above samples were analyzed to confirm which substances were present or absent on the fiber surface, and that the sheath of the bicomponent fibers contained a blend of PCL and dECM.

#### Immunocytochemical staining

Immunocytochemical staining was performed to qualitatively assess the presence of ECM proteins in the PU-PCL/dECM fibers. PU-PCL samples served as the negative control. Two specimens from each sample type were stained to detect collagen I, III, elastin, laminin, and fibronectin. **Table S1** provides information on the primary and secondary antibodies used for staining. The samples were permeabilized with 0.1% Tween-20 and 0.1% Triton X-100. Next, the samples were incubated with a blocking buffer solution (2% bovine albumin serum + 2% animal serum in PBS+0.1% tween-20) for 60 minutes to decrease nonspecific antibody binding, followed by overnight incubation of primary antibodies at 4 ^°^C. After incubation, the specimens were washed in PBS + 0.1% Tween-20 for 5 minutes three times. Secondary antibodies were then diluted at relevant concentrations in 10% blocking buffer + 90% PBS, as specified in Table 3.2, and incubated for 60 minutes. Imaging was performed using a fluorescent microscope (EVOS FL Auto 2, Invitrogen).

### 5.6. Characterization of the surface hydrophilicity of the three electrospun nanofiber types

Contact angle measurements were taken to evaluate the hydrophilicity of each sample type. 4 specimens of 2.54 cm x 2.54 cm were cut from each sample type. 3 µl water droplets were suspended on the top (the surface on which fibers got deposited continuously) and bottom (the surface that touches the copper shim) surfaces of each specimen and 12 contact angle measurements were taken per side per specimen using a goniometer (Dataphysics OCA System, FDS Corp). Using those readings, the average contact angles were calculated for each sample type.

### 5.7. Characterization of the mechanical properties of the three electrospun nanofiber types

The mechanical properties of the PU-dECM/PCL fiber samples and controls samples; PU-PCL and dECM/PCL were evaluated using Suh et al.’s modified tensile test adapted from ASTM standard D5035, “Textile Strip Method”^[52]^. For each sample type, 5 specimens with dimensions 4 x 1 cm^2^ were tested and the respective stress-strain graphs were obtained using a tensile tester (MTS Criterion^TM^ Model 43). A gauge length of 1 inch (2.54cm) of each specimen was tested using a load cell of 5N, at a 0.01 mm/s strain rate and break sensitivity of 50%. Using the data from the linear region of the stress-strain curves, Young’s modulus for each specimen was calculated and finally, the average was taken for each sample type. For easy handling of the samples, they were attached to C-cards that were made using 3” x 5” index cards, using thin double-sided tape, and lab tape.

### 5.8. Analysis of the enzymatic degradation of the three electrospun nanofiber types by weight

The degradation profiles of the three fiber types were studied by maintaining 50 specimens of 0.5 x 0.5 cm^2^ from each type in an enzyme solution prepared by mixing equal volumes of 0.25mg/ml collagenase – Type I (Fisher) in 0.1M Tris-HCl and 0.005M CaCl_2_ and 0.0025 mg/ml Pseudomonas lipase (Type XIII, ≥15 units/mg solid, Millipore Sigma) in PBS as reported by Suh et al^[52]^. Before immersing in the said enzyme mixture, the samples were dried in a desiccator for 24 hours and the dry weight of each sample was measured and recorded. Then they were placed in 12-well plates (specimen per well) and 1 ml of the enzyme solution was added to each well. The plates with specimens were incubated at 37° C for 4 weeks and the change in mass and morphology with time were studied. On days 1, 7, 14, 21, and 28 10 specimens from each sample type were taken out of the enzyme solution and dried in a desiccator overnight. On the respective following days (days 2, 8, 15, 22, and 29) the dry masses were measured. Using those readings, the average percent change in mass was calculated for each sample type at each time point using the following equation.

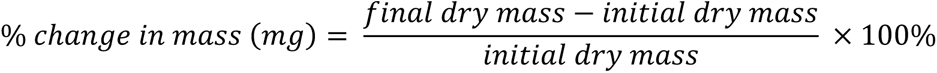

To observe the structural changes in the fibers, SEM images of specimens belonging to different sample types were obtained using a scanning electron microscope (Hitachi TM4000) with 15kV accelerating voltage and BSE (backscattered electron) mode, under x2500 magnification. 2 specimens per sample type were used for this and they were sputter coated with gold/palladium prior to imaging.

### 5.9. Biocompatibility analysis using induced pluripotent stem cell (iPSC) derived cardiac differentiation

iPSCs were acquired from WiCell (19-9-7T) and cultured in 6-well dishes coated with LN-521 (BioLamina) overnight. Cells were maintained in Essential 8 medium (Gibco) and subcultured at 90% confluence using ReLeSR (Stem Cell Technologies) as a dissociating agent, following the manufacturer’s protocol. For differentiation, a previously established protocol was followed.^[30]^ Briefly, the cells were transferred to 12-well dishes and cultured in Essential 8 medium until 99-100% confluence. The medium was then switched to RPMI 1640 medium (Gibco) with B27 supplement minus insulin and containing 6µM CHIR99021 (Tocris) for 24 hours. Afterward, the medium was changed to RPMI 1640 with B27 supplement minus insulin for an additional 48 hours. WNT-inhibition was achieved using IWR1 (Tocris) at 12µM concentration, mixed with a 1:1 ratio of spent medium for 24 hours, followed by RPMI 1640 with B27 supplement minus insulin. RPMI 1640 supplemented with B27 was then used, with media changes occurring every Monday, Wednesday, and Friday until contractile cells were obtained.

Once contractility was achieved, the cells were dissociated using ReLeSR, seeded onto sterilized samples, and maintained in RPMI 1640 supplemented with B27 and 20% fetal bovine serum. The samples were then assessed using LIVE/DEAD™ (Invitrogen) at days 1, 3, 5, and 7 of seeding, according to the manufacturer’s protocol. CM-specific staining was performed using the above IFF/ICC protocol. PCR was performed to relatively quantify the expressions of cardiac troponin T, alpha actinin, and myosin heavy chain 7, where iPSC-CM on TCPS was used as a control.

### 5.10. Calcium Imaging of iPSC-CMs

iPSC-CMs were cultured on TCPS for 9 days until spontaneous contractile activity was observed. Following the onset of contraction, cells were enzymatically dissociated and seeded onto the experimental scaffold samples. Live-cell calcium imaging was performed using Fluo-8 AM (VWR, Cat# 76483) dissolved in Pluronic F-127 (VWR, Cat# P3000MP) and 2.5 mM probenecid (VWR, Cat# P36400) in Tyrode’s buffer. Culture media were aspirated and replaced with the staining solution, followed by incubation at 37 °C for 45 minutes. After staining, samples were washed three times with Tyrode’s buffer and incubated again in fresh Tyrode’s buffer at 37 °C for an additional 45 minutes to allow for de-esterification of the dye.

Fluorescence imaging was conducted using the GFP channel on the EVOS FL Auto 2 system, capturing time-lapse images at 20 frames per second (FPS). Calcium transients were analyzed using the *Time Series Analyzer* plugin in ImageJ (FIJI). Fluorescence intensity was normalized to ΔF/F₀ = (F − F₀)/F₀, where F₀ represents baseline fluorescence and F the peak fluorescence intensity. Inter-beat interval (IBI) and calcium decay time (T_50_) were calculated manually based on the time differences between fluorescence peaks and the time to 50% decay from the peak, respectively.

### 5.11. Statistical analysis

The results of all the quantitative assessments were analyzed using GraphPad software and expressed as mean ± standard error of the mean. Statistical significance was evaluated using one-way and two-way analysis of variance (ANOVA). Differences among means with p < 0.05 were considered statistically significant and were displayed as \**p* < 0.05, \*\**p* < 0.01, and \*\*\**p* < 0.001.

## Acknowledgements

The schematics and TOC graphic were made using BioRender. This work was performed in part at the Analytical Instrumentation Facility (AIF) at North Carolina State University, which is supported by the State of North Carolina and the National Science Foundation (award number ECCS-2025064). The AIF is a member of the North Carolina Research Triangle Nanotechnology Network (RTNN), a site in the National Nanotechnology Coordinated Infrastructure (NNCI).

## Supporting Information

Supporting Information is available online or from the corresponding author.

