## Supplemental data for "Coaxially Electrospun Myocardial dECM- based Nanofibrous Scaffolds Demonstrate Enhanced Cardiomyocyte Adhesion and Function"

**Materials for electrospinning**

1,1,1,3,3,3-hexafluoro-2-propanol (HFIP) (Thermo Fisher), Polycaprolactone (PCL) (Mw 150000, Scientific Polymer Products, Inc.), Polyurethane Selectophore$^{TM}$ (PU) (Mw 100 kDa, Sigma-Aldrich), coaxial needle (17G-22G), normal needle (18G), syringes (Fisherbrand 10ml), copper 110 shim stock (Trinity Brand Industries), tube (McMaster-Carr), 1-channel syringe pump (New Era Pump Systems), voltage supply (Gamma High Voltage Research, Inc).

**FIGURES**

**Figure S1. Visual observation of the relative abundance of the fibers with core-sheath geometry.** Cross-sectional TEM Images of coaxially electrospun PU-dECM/PCL fibers at (a) 10k magnification, scale bar = 2 μm (b) 25k magnification, scale bar = 1 μm. Inner white dashed lines indicate the border of the core, while the outer yellow lines indicate the border of the sheath.

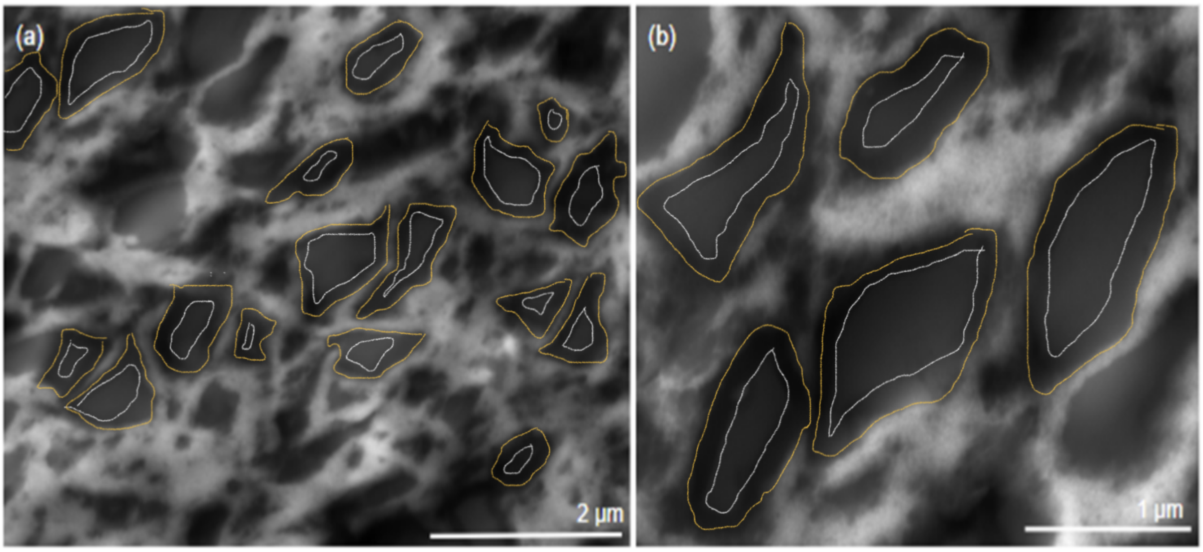


**Figure S2. Surface hydrophilicity of PU-dECM/PCL, dECM/PCL, and PU-PCL fibers.** Goniometer measurements of the contact angles of PU-dECM/PCL samples (a) top (b) bottom, dECM/ PCL samples (c) top (d) bottom, PU-PCL samples (e) top (f) bottom

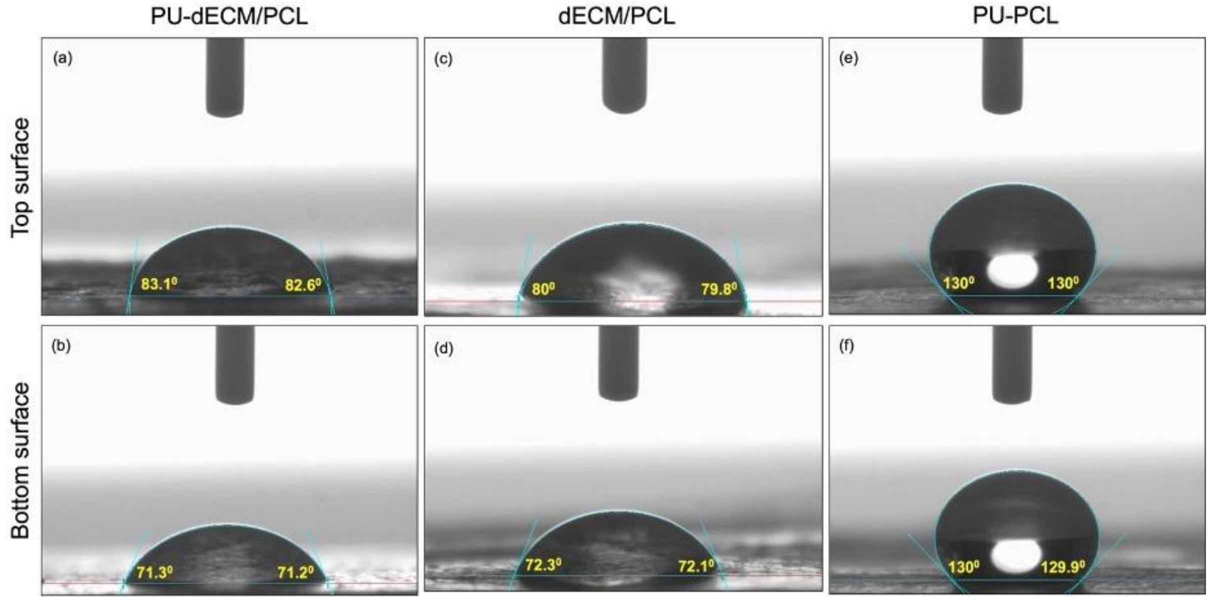


**Figure S3. Optimization of PU-PCL fiber fabrication parameters.** SEM images of the PU-PCL fibers fabricated using different voltages, obtained at x500, scale bar = 50µm

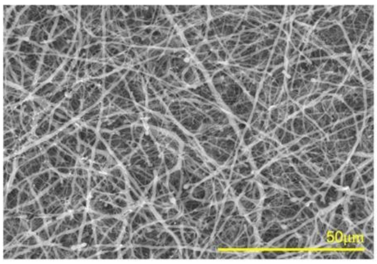


Positive voltage: 20 kV
Negative voltage: 10 kV

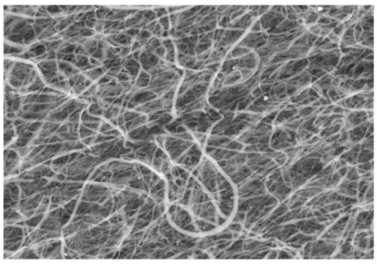

Positive voltage: 20 kV
Negative voltage: 0 kV

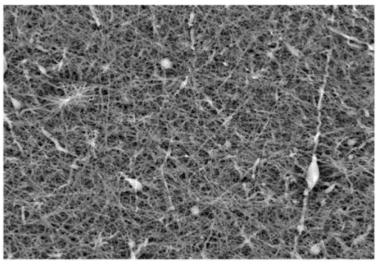

Positive voltage: 15 kV
Negative voltage: 0 kV

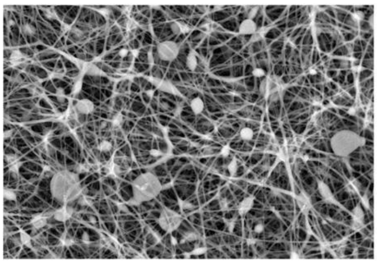

Positive voltage: 12 kV
Negative voltage: 0 kV

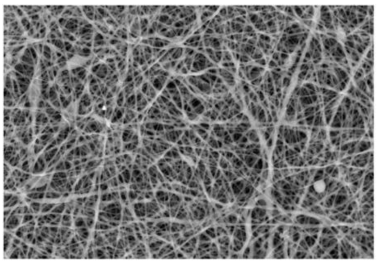

Positive voltage: 10 kV
Negative voltage: 0 kV

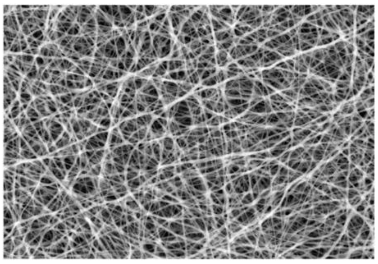

Positive voltage: 8 kV
Negative voltage: 0 kV

### TABLES

**Table S1.** Primary and secondary antibodies used to stain the PU-dECM/PCL and PU-PCL nanofibers to detect the presence of ECM proteins collagen I, III, elastin, laminin, and fibronectin

| | Collagen I | Collagen III | Elastin | Laminin | Fibronectin |
|---|---|---|---|---|---|
| Primary antibody | collagen I monoclonal antibody | anti-collagen III antibody | anti-elastin antibody | Laminin Rat anti-Human, Mouse, Porcine, Clone: A5, MUbio | Fibronectin Antibody (A-11) |
| Vendor | Fisher | Abcam | Abcam | Nordic | Abcam |
| Catalog number | MA1-26771 | ab7778 | ab21610 | 50-210-1098 | ab2413 |
| antibody: blocking buffer (volume ratio) | 1:100 | 1:200 | 1:100 | 1:100 | 1:500 |
| Secondary antibody | IgG1 (y1) Goat anti-mouse alexa fluor 488 | IgG (H+L) cross adsorbed goat anti-rabbit alexa fluor 594 | IgG (H+L) cross adsorbed goat anti-rabbit alexa fluor 488 | Donkey anti-Rabbit IgG (H+L) Highly Cross-Adsorbed Secondary Antibody, Alexa Fluor™ 568 | Goat anti-Rabbit IgG (H+L) Cross-Adsorbed Secondary Antibody, Alexa Fluor™ 350 |
| Vendor | Invitrogen™ | Fisher | Fisher | Fisher | Fisher |
| Catalog number | A21121 | A11012 | A11008 | A78946 | A11046 |
| antibody: blocking buffer (volume ratio) | 1:100 | 1:100 | 1:100 | 1:100 | 1:100 |

**Table S2:** RTqPCR primers used for the comparison of genetic expression of iPSC-CMs on nanofibrous scaffolds.

| Primer | Vendor | Forward Sequence | Reverse Sequence |
|---|---|---|---|
| MYH7 | IDT | CCT GAG GGA CAG TTC ATT GAT AG | GCC ATC TCC TTC TCT CTT TCT G |
| ACTN2 | IDT | CTA TGC CTC TGG ACG CAC AAC T | CAG ATC CAG ACG CAT GAT GGC A |
| TNNT2 | Qiagen | Quantitect Assay: TNNT2 (QT00089782) | |
| GAPDH | IDT | ACC ACA GTC CAT GCC ATC AC | TCC ACC ACC CTG TTG CTG TA |

**Table S3:** Primary antibodies for cardiac staining via IFF/ICC

| Antibody | Vendor/catalog# |
|---|---|
| Cardiac Troponin T | Developmental Studies Hybridoma Bank (DSHB)/RVC2 |
| Alpha Actinin | ThermoFisher/ MA1-22863 |
